# Microenvironment-informed inference of transcriptional progression geometry

**DOI:** 10.64898/2026.08.21.746284

**Authors:** Seibi Kobara, Sheikh A. Rahman, Susan P. Ribeiro, Craig M. Coopersmith, Rishikesan Kamaleswaran

## Abstract

We present BIOCURRENT, a causal inference framework that reconstructs donor-specific pseudotime geometry in transcriptomic data. By modeling gene expression as a function of baseline characteristics, microenvironmental context, and latent pseudotime, BIOCURRENT enables comparison of compressed or expanded progression intervals across transcriptional state transitions. We introduce ΔΔ*T*, a geometry-based estimator that quantifies differences in pseudotime intervals across conditions, enabling evaluation of changes in pseudotime intervals under hypothetical modulation of microenvironmental programs. Applications to thymic T-cell developmental lineages and to COVID-19 immune dysregulation reveal condition- and donor-specific distortions of progression intervals. Counterfactual simulation links microenvironmental context to changes in specific intracellular state transition intervals. By localizing deviations in pseudotime geometry, BIOCURRENT identifies whether shifts in transcriptomic programs emerge early or later along transcriptomic coordinates and reveals upstream programs associated with these distortions. Such localization supports transcriptional stage-aware mechanistic hypotheses and suggests candidate intervention checkpoints in complex biological systems.

## INTRODUCTION

Advanced transcriptomic technologies, including single-cell RNA sequencing (scRNA-seq) and spatial transcriptomics, have enabled high-resolution characterization of cellular states and their organization within tissues, substantially transforming our understanding of biological systems and informing therapeutic discovery^1^. By resolving heterogeneous cell populations and intracellular molecular programs, these approaches provide a foundation for studying dynamic biological processes such as development, immune activation, and disease progression^2,3^.

Despite substantial advances in molecular profiling, many diseases and pathological conditions with complex biology still lack therapies that reliably target the appropriate biological processes at the right stage of pathology, reflecting an incomplete understanding of the timing and dynamics of cellular state transitions. For example, in critical illnesses such as sepsis, which remains a major contributor to global mortality^4–6^, therapeutic strategies are largely supportive, underscoring the difficulty of translating molecular insights into effective, stage-specific interventions^7^. More broadly, the precise molecular mechanisms underlying pathological progression, as well as the identification of actionable intervention targets, optimal dosing strategies, and critical time windows for intervention, remain incompletely understood. Addressing these challenges requires analytical frameworks capable of capturing not only static cellular heterogeneity but also the individualized patterns of biological progression underlying complex pathological conditions.

To characterize such dynamic processes, numerous bioinformatic methods have been developed to infer a latent pseudotime^8^, a low-dimensional transcriptional coordinate intended to approximate biological progression along a cell-state trajectory. Existing approaches span a wide range of modeling assumptions, from linear trajectory inference with fixed prior structure to complex branching and multi-lineage reconstructions. Early methods addressed non-linear cell-state transitions using manifold learning techniques, such as diffusion-based pseudotime inference^9^, locally linear embedding-based approaches^10^, and a Gaussian process-derived latent variable model^11^, enabling reconstruction of smooth developmental progression. Graph- and tree-based methods further extended trajectory inference to complex topologies by modeling cell-state relationships using minimum spanning trees^12^ or neighborhood graphs^13^, allowing identification of branching and bifurcation events. Additional probabilistic frameworks have been proposed to capture uncertainty in lineage inference^14,15^.

More recently, advances in deep learning and large-language models have enabled increasingly expressive models for dynamic cell-state inference, including approaches that model stochastic transitions and responses to environmental perturbations^16,17^. Collectively, these methods have substantially expanded the scope of trajectory analysis in single-cell transcriptomics. Nevertheless, most existing frameworks are developed under implicit assumptions of a homogeneous biological context, relying on controlled experimental simulation and data. This assumption likely does not hold in complex disease conditions, where observed transcriptomic profiles and cell-state transitions reflect heterogeneous baseline characteristics and diverse tissue microenvironment, complicating interpretation of inferred pseudotime as a comparable measure of biological progression.

Conventionally, pseudotime reconstruction is achieved by learning a latent progression co-ordinate directly from observed transcriptomic expression measured by scRNA-seq or spatial transcriptomics (**Figure S1**). This modeling paradigm is well-suited for descriptive reconstruction of transcriptional continua within a homogeneous context. However, conceptual limitations arise when pseudotime is interpreted as a proxy for biological progression in downstream analyses, such as modeling gene expression as a function of pseudotime to identify progression-associated gene signatures or to compare trajectories across experimental or clinical phenotypes. Because pseudotime is inferred solely from gene expression, its interpretation as a biological progression implicitly assumes that the mapping between expression and biological progression is invariant across samples, conditions, and tissue environments.

In practice, baseline characteristics and microenvironmental factors—including circulating cytokines and ligands, tissue architecture, and inflammatory state—can modulate both underlying biological progression and gene expression programs^18–20^. When pseudotime is inferred directly from gene expression, these ancestry influences can distort estimation of trajectory geometry, and apparent differences in pseudotime-associated genes or trajectory structure may be difficult to attribute to intrinsic changes in cell-state progression or context-dependent environmental programs. In addition, the same amount of biological progression may correspond to different pseudotime intervals across donors or conditions. For example, donors with heightened proliferative or activation states may exhibit compressed pseudotime scales. Conventional pseudotime models do not explicitly account for such donor-specific scaling effects. Together, these limitations obscure the interpretation of pseudotime-associated gene programs and complicate the identification of mechanistic targets for therapeutic intervention.

To address this problem, we have developed a computational method, named **B**ayesian-**I**nf**O**rmed **C**a**U**sal **R**egulatory **R**emodeling of **EN**vironment-conditioned cellular **T**ransition (BIOCURRENT) (**Figure 1**). BIOCURRENT models gene expression as a function of latent biological progression together with sequence-related covariates, baseline characteristics, and microenvironmental inputs, such as circulating ligands and tissue architectural features, enabling joint estimation of intrinsic cellular transcriptional dynamics and context-dependent regulatory effects. By incorporating donor- and state-level random effects in pseudotime priors, BIOCURRENT supports identification of donor-specific pseudotime coordinates, comparative analysis of trajectory geometry across conditions, and counterfactual simulation of microenvironmental perturbations on cellular states. We evaluated BIOCURRENT using synthetically generated multi-donor scRNA-seq data and time-stamped transcriptomic data^21^ to assess recovery of true temporal coordinates. We further deployed BIOCURRENT on spatial transcriptomic data of thymic T cell developmental processes^18^ and naive CD4 T cell activation dynamics in Coronavirus Disease 2019 (COVID-19)^2^.

**Figure 1.**
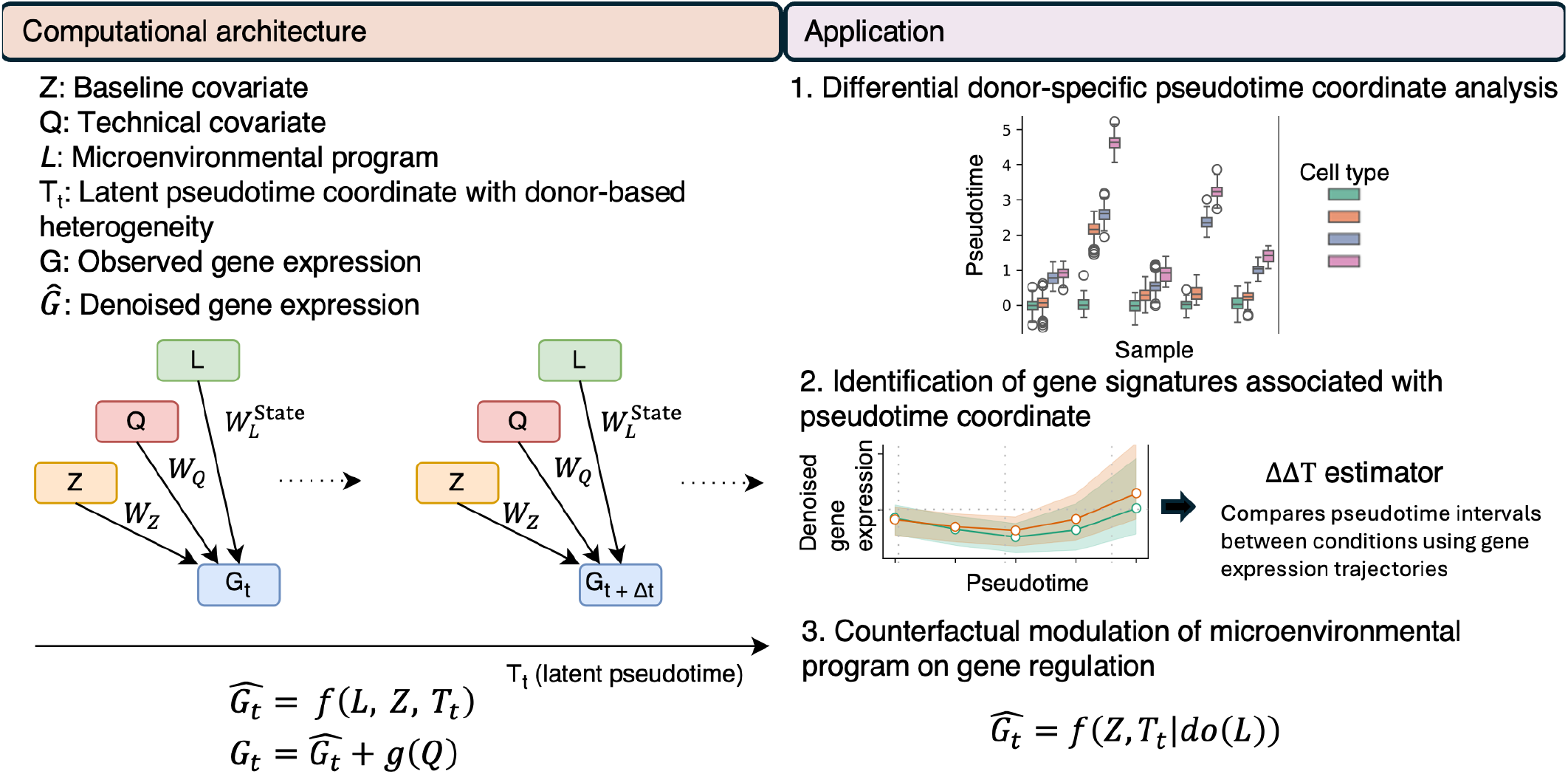
Overview of BIOCURENT. BIOCURRENT (**B**ayesian-**I**nf**O**rmed **C**a**U**sal **R**egulatory **R**emodeling of **EN**vironment-conditioned cellular **T**ransition) is a causal inference framework for transcriptomic pseudotime analysis that estimates donor-specific latent pseudotime coordinates while conditioning on baseline covariates (*Z*), sequence-related technical factors (*Q*), and microenvironmental context (*L*). By jointly modeling these components, BIOCURRENT produces denoised gene expression estimates that are deconfounded from technical effects (left panel). The right panel illustrates three key analytical capabilities of BIOCURRENT: (1) estimation of donor- and condition-specific pseudotime geometry; (2) identification of genes associated with expansion or compression of pseudotime intervals across defined transcriptional state transitions; and (3) counterfactual simulation of microenvironmental perturbations to infer their effects on stage-specific transcriptional dynamics.

## RESULTS

### Probabilistic model design and synthetic data experiments

We first evaluated BIOCURRENT using synthetic data generated directly from the proposed probabilistic model (see Method details). Synthetic datasets were constructed with predefined baseline characteristics, microenvironmental signatures, sequence-related technical covariates of scRNA-seq, and pseudotime coordinates. We systematically varied key parameters, including the number of linear transcriptomic states, the number of cells per state, and the sparsity of microenvironmental coefficients (fraction of near-zero values), which parameterize the effect of microenvironmental programs on gene expression.

Pseudotime coordinates were modeled using ordered latent progression coordinates with Gaussian priors, allowing cells within each transcriptomic state to vary continuously along pseudotime. All latent nodes were simulated from Gaussian distributions with standard deviation *σ* = 0.5. Independent noise, modeled as Gaussian with *σ* = 0.75, was added to gene expression to reflect technical and biological variability. For each simulation scenario, 50 independent synthetic datasets were generated.

BIOCURRENT accurately recovered latent pseudotime across a range of simulation settings. As shown in **Figure 2A**, the correlation between true and inferred pseudotime reached 0.920 (95% confidence interval [CI]: 0.903, 0.935) when simulating four linear cell state transitions with the mean of 100 cells per transcriptomic state and 95% sparsity of microenvironment coefficients. Increasing the number of transcriptomic states to eight showed stable model performance (correlation coefficient: 0.966, 95% CI: 0.957, 0.973). Overall, BIOCURRENT robustly recovered underlying pseudotime despite high levels of noise (Gaussian noise with *σ* = 0.75) in gene expression.

**Figure 2.**
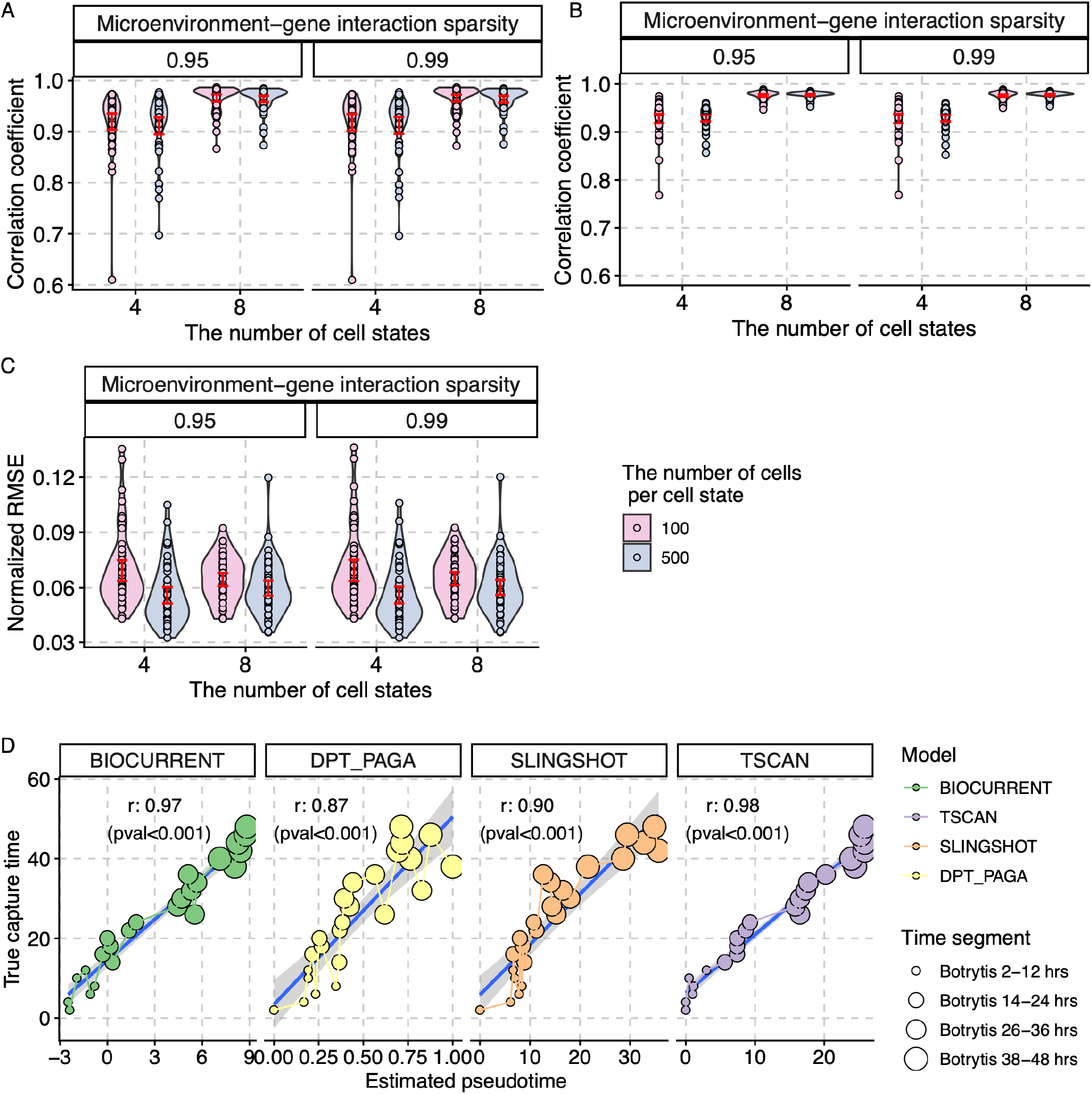
Synthetic data experiments and validation using time-stamped data. (A) Accuracy of pseudotime recovery, measured by Pearson correlation coefficients, across simulation settings varying the numbers of transcriptional states, the number of cells per state, and the sparsity of microenvironment coefficients (fraction of near-zero values). Red vertical lines indicate Wald 95% confidence intervals (CIs). (B) Accuracy of donor-specific pseudotime estimation, measured by Pearson correlation coefficients, across five simulated donors under the same simulation parameter combinations. Red vertical lines indicate Wald 95% CIs. (C) Accuracy of denoised gene expression estimation, measured by normalized root mean squared error (RMSE). Denoised gene expression is defined as expression after the removal of sequence-related covariate effects. Red vertical lines indicate Wald 95% CIs. (D) Benchmarking of BIOCURRENT against existing pseudotime methods using time-stamped transcriptomic data. Accuracy of clock-time recovery was measured by Pearson correlation coefficients between true clock time and estimated pseudotime.

To assess donor-specific pseudotime identification, correlation coefficients between true and inferred pseudotime were computed separately for each donor and summarized across simulations. The mean donor-level correlation was 0.928 (95% CI: 0.918, 0.938) when simulating four linear cell state transitions with the mean of 100 cells per transcriptomic state and 95% sparsity of microenvironment coefficients (**Figure 2B**), showing reliable recovery of donor-specific progression coordinates.

Identification of denoised gene expression, defined by eliminating the effect of sequence-related covariates, was evaluated using root mean squared error divided by the true standard deviation (normalized RMSE). In simulation under four linear cell state transitions with the mean of 100 cells per transcriptomic state and 95% sparsity of microenvironment coefficients, the normalized RMSE was 0.069 (95%CI: 0.063, 0.075) (**Figure 2C**).

### Benchmarking pseudotime inference using time-stamped transcriptomic data

We next assessed the biological identification performance of BIOCURRENT using bulk RNA-seq data with known time stamps. BIOCURRENT was compared against nearest–neighbor–based diffusion-based, and tree-based algorithms, including Slingshot, Diffusion Pseudotime (DPT), and TSCAN. Preprocessing steps and analytical settings are provided in Method details. Briefly, we used publicly available transcriptomic data of a *Arabidopsis thaliana* leaf following *Botrytis cinerea* infection^21^. Twenty-four time-stamped gene expression profiles were grouped into four ordered biological states. To evaluate recovery of temporal ordering, the biological clock time was masked for six profiles within each state group. State group assignments and their ordering were provided to each method, and inferred pseudotime coordinates were compared against the withheld time stamps. BIOCURRENT achieved a correlation of 0.97 (p-value *<* 0.001) between inferred pseudotime and true time stamps, ranking second among the evaluated methods (**Figure 2D**).

### Reconstruction of T cell development in thymic cortical regions

We first applied BIOCURRENT to publicly available spatial transcriptomics of fetal and postnatal thymic samples^18^. Thymic T cell development represents a biologically regulated process with a well-characterized progression structure, providing a biologically constrained setting to evaluate pseudotime geometry inference. The original study identified a conserved T cell receptor (TCR) signaling across fetal and postnatal samples, with enrichment of cycling and proliferative gene programs in fetal thymus^18,22^. We extend this analysis by testing whether BIOCURRENT-inferred pseudotime coordinates differed between fetal and postnatal samples.

We modeled five T cell developmental states—early T cell progenitors (ETP), double-negative (DN) early, DN late, double-positive (DP) early, and DP late. Baseline covariates included biological sex and normalized gestational age centered on 40 weeks post-conception. To mitigate sample- and transcriptional state–imbalance, we applied inverse-frequency weighting during model fitting, ensuring that underrepresented samples and states contributed comparably to the optimization objective (see Method details). Estimated pseudotime coordinates were summarized within each developmental state, and pseudotime intervals between adjacent developmental states were compared between fetal and postnatal samples using Welch’s t-test. Detailed gene expression decomposition using cell2location^23^, preprocessing, quality control procedures, and data preparation are provided in Method details.

The pseudotime intervals from DN early to DN late and from DN late to DP early were significantly shorter in fetal samples (p-value=0.0486, 0.0447, respectively) (**Figure 3A** and **Figure S2**) with a spatial pseudotime projection (**Figure 3B** and **Figure S3**). Statistical tests were performed using R 4.4.1, and spatial visualization was performed using Squidpy^24^.

**Figure 3.**
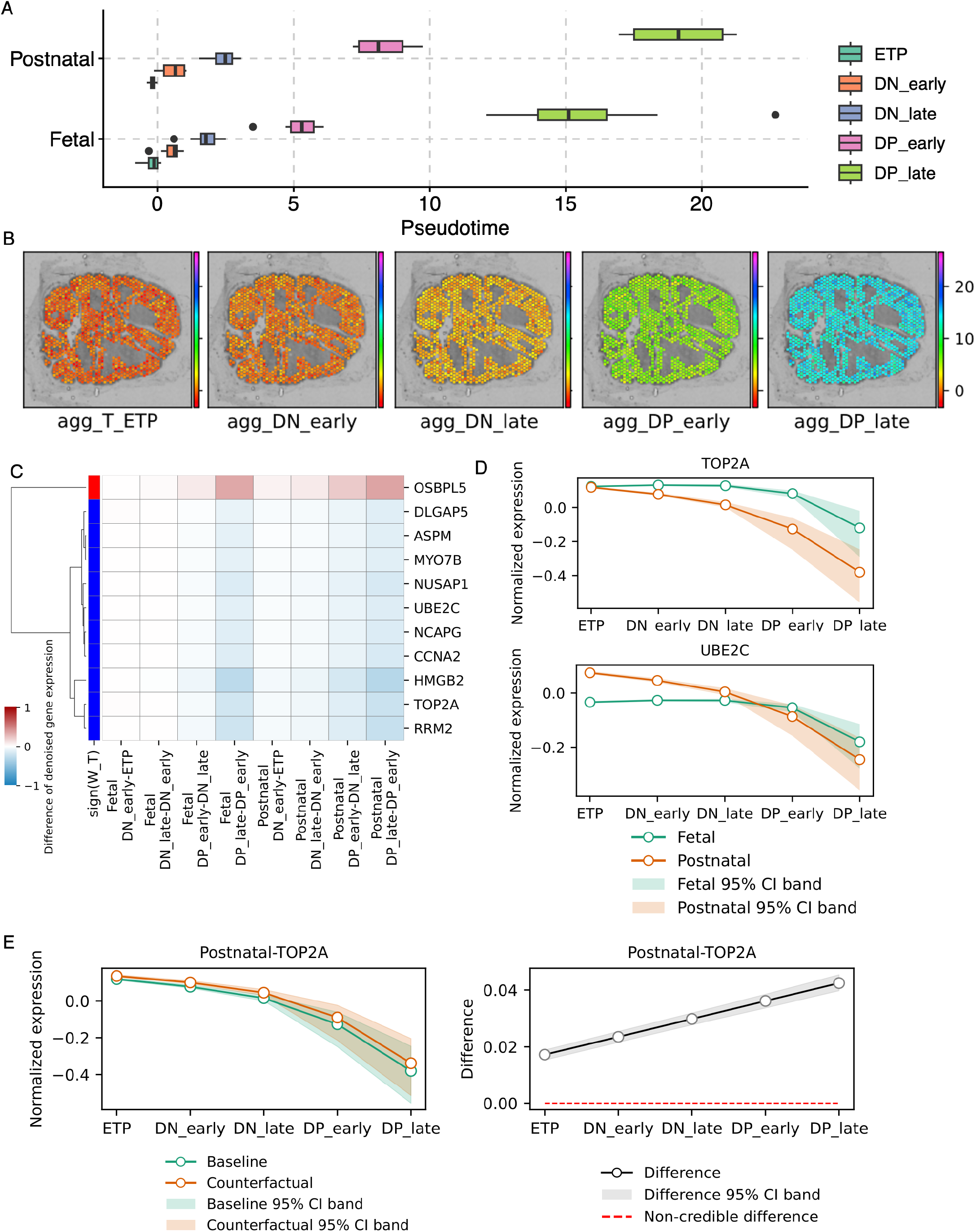
Pseudotime estimation in fetal and postnatal thymic spatial transcriptomics. (A) Posterior median of donor-specific pseudotime coordinates in five T cell developmental lineages for fetal and postnatal samples. Pseudotime estimates are summarized by the median and interquartile range for each transcriptional state. (B) Spatial projection of inferred posterior median of pseudotime across T cell developmental lineages in the thymic cortex (representative fetal sample shown). (C) Differences of pseudotime-associated genes based on 95% credible intervals (CrIs) across transcriptional state transitions with corresponding signs of pseudotime coefficients (*W*_*T*_) that link between gene expression and pseudotime. (D) Estimated denoised gene expression across T cell developmental lineages for fetal and post-natal samples. Shaded regions represent 95% CrIs. (E) The left panel shows counterfactual denoised gene expression under a counterfactual intervention in which the non-negative matrix factorization (NMF) module 3 of postnatal samples was reduced to the mean level of NMF module 3 observed in prenatal samples. The right panel shows the difference between counterfactual- and baseline-denoised gene expression. Shaded regions represent 95% CrIs. The red dashed line represents a difference of 0.

### Genes associated with narrowed pseudotime during fetal thymic development

To investigate molecular programs associated with the shortened pseudotime interval in fetal compared with postnatal samples, we identified credible genes that were associated with narrowed pseudotime across DN early to DN late and DN late to DP early transitions in fetal samples.

In BIOCURRENT, differences in pseudotime interval across phenotypes are modeled through the combination of 1) gene-specific pseudotime coefficients and 2) denoised gene expression changes across state transitions. Formal derivation is provided in Method details. Using this framework, 11 of 500 genes were credibly associated with the narrowed pseudotime interval in fetal samples relative to postnatal samples based on 95% credible intervals (CrIs) (**Figure 3C**). These genes were cell-cycle and proliferation-related programs, including DNA Topoisomerase II Alpha (TOP2A) and Ubiquitin Conjugating Enzyme E2 C (UBE2C). A heatmap with hierarchical clustering was generated using Seaborn^25^.

TOP2A expressions were comparable between fetal and postnatal samples at the ETP state (**Figure 3D** and **Figure S4**). However, their trajectories differed substantially thereafter: fetal samples exhibited relatively stable TOP2A expression across subsequent development states, whereas postnatal samples showed a progressive decrease. A similar pattern was observed in UBE2C, which remained stable from ETP to DP early states in fetal thymus but declined across these states in postnatal samples.

### Cortical epithelial microenvironment

BIOCURRENT is designed to model modulatory effects of the tissue microenvironment on intra-cellular gene regulatory programs. In the context of thymic development, we obtained cortical thymic epithelial cell (cTEC)-specific gene expression for each spatial spot using cell2location. Estimated cTEC-derived gene expression profiles were then conditioned to infer intracellular T cell developmental state transitions.

To represent the microenvironment, we first obtained 1,000 highly variable genes across fetal and postnatal samples. Among these genes, we explicitly modeled expression of the key cTEC-derived ligands (DLL4, CXCL12, CCL25, and WNT4). The remaining 996 highly variable cTEC-associated genes were summarized into five modules using non-negative matrix factorization (NMF). Ranked gene sets based on NMF were used to perform gene-set-enrichment analysis (GSEA) using the Gseapy package^26^ to annotate each representative immune program (**Table S3**. The top contributing genes for each module are provided in **Figure S5A**.

Briefly, NMF0 captured peptidase activity program (e.g., PRSS16 and PSMB11). NMF1 represented epithelial and structural development (e.g., KRT5 and DSP). NMF2 represented antigen and peptide processing capacity (e.g., CTSV and PITHD1). NMF3 represented apoptotic stress signal (e.g., GAS6 and NUPR1). Lastly, NMF4 represented membrane lipid remodeling (e.g., PLTP and PASK). Descriptive comparisons of observed gene expression and microenvironmental feature values between fetal and postnatal samples are shown in **Figure S5B**. Estimated associations between pseudotime-linked genes and microenvironmental features are provided in **Figure S5C** as part of the BIOCURRENT framework.

### Counterfactual analysis of microenvironmental contributions to pseudo-time compression

To explore microenvironmental features that may contribute to fetal proliferative characteristics in DN early to DN late and DN late to DP early transitions, we performed counterfactual simulations within the BIOCURRENT framework. In the counterfactual analysis, microenvironmental variables in postnatal samples were altered, targeting the mean values of fetal samples while holding other parameters fixed, including inferred pseudotime coordinates. This design estimates the conditional effect of microenvironmental normalization on pseudotime contrast without assuming restoration of pseudotime positions.

We defined a ΔΔ*T* estimator, representing the difference in inferred pseudotime intervals between phenotypes, for example, the different pseudotime interval from DN early to DN late between fetal and postnatal samples. ΔΔ*T* was subsequently used to quantify the counterfactual normalization of the altered pseudotime intervals between phenotypes.

Counterfactual reduction of apoptotic stress signal (NMF3) in postnatal samples was causally associated with a reduction of ΔΔ*T* (-0.465 [95%CrI: -0.477, -0.448]) for the interval from DN early to DN late, reducing the differences in pseudotime intervals between fetal and postnatal samples. Similarly, counterfactual increases in DLL4 and NMF1 (epithelial and structural development) in postnatal samples were causally associated with decreases in ΔΔ*T* of -0.141 [95%CrI: -0.144, -0.138] and -0.636 [95%CrI: -0.647, -0.626], respectively, compared with baseline, indicating partial alleviation of pseudotime intervals between DN early and DN late across fetal and postnatal samples.

At the gene level, reduction of NMF3 in postnatal led to a credible increase in denoised expression of TOP2A based on 95% CrIs (**Figure 3E**). In contrast, counterfactual intervention of DLL4 or NMF1 resulted in minimal changes in TOP2A expression (**Figure S6**).

Together, these results demonstrate that BIOCURRENT localizes distinct pseudotime transition geometries between fetal and postnatal thymic T-cell developmental trajectories. Counter-factual analyses indicate that higher expression of DLL4 and structural-associated signatures, as well as lower stress-associated signatures in cTECs, are each associated with expanded proliferative transition intervals during early T-cell development.

### BIOCURRENT application to scRNA COVID-19 dataset

For our second real-data application of BIOCURRENT, we analyzed scRNA-seq data from the COVID-19 multi-omics blood atlas (COMBAT) consortium^2^. The original COMBAT study investigated all-cause sepsis excluding SARS-CoV-2 infection, as well as three discretized severity categories of COVID-19 based on World Health Organization (WHO) guidelines: mild cases (no requirement for supplemental oxygen), severe cases (oxygen saturation SaO_2_ ≤ 93% on air but not requiring mechanical ventilation), and critical cases (requiring mechanical ventilation). This study demonstrated that COVID-19 severity is associated with coordinated dysregulation of innate and adaptive immunity, including impaired antigen presentation in innate immune cells and altered cell cycle and redox state pathways in adaptive immune cells. We hypothesized that the transcriptional landscape of adaptive immune cells, as captured by pseudotime geometry, varies across COVID severity, and distinct microenvironment contexts contribute to impaired adaptive immune response across severity categories. To test this hypothesis, we focused on CD4 T cell transitions from naive to activated states. Specifically, we defined three transcriptional states along with this continuum: naive (level 0), effector (level 1), and terminal (level 2). The corresponding cell-type annotations from the original COMBAT study are provided in **Supplementary Table S2**.

Baseline characteristics included age, sex, pre-existing comorbidities, and other variables as described in the Methods. Microenvironmental features were defined using ligand gene expression and inflammatory state signatures derived from classical monocytes, non-classical monocytes, dendritic cells, and natural killer cells. BIOCURRENT was then used to estimate pseudotime coordinates across mild COVID-19, severe COVID-19, critical COVID-19, and sepsis patient groups. As in the thymic analysis, inverse-frequency weighting was applied to mitigate sample- and transcriptional state–imbalance (see Method details).

### Pseudotime compression in critical COVID-19 CD4 T cells

Estimated pseudotime intervals over transcriptional states were compared across COVID-19 severity and sepsis using Welch’s one-way ANOVA, followed by pairwise comparisons using the Games–Howell post hoc test. We observed a statistically significant narrowing of pseudotime coordinates between the naive (level 0) and effector (level 1) states in critical COVID-19 samples compared with other groups (adjusted p value *<* 0.05) (**Figure 4A and Figure S7C, D, and E**).

**Figure 4.**
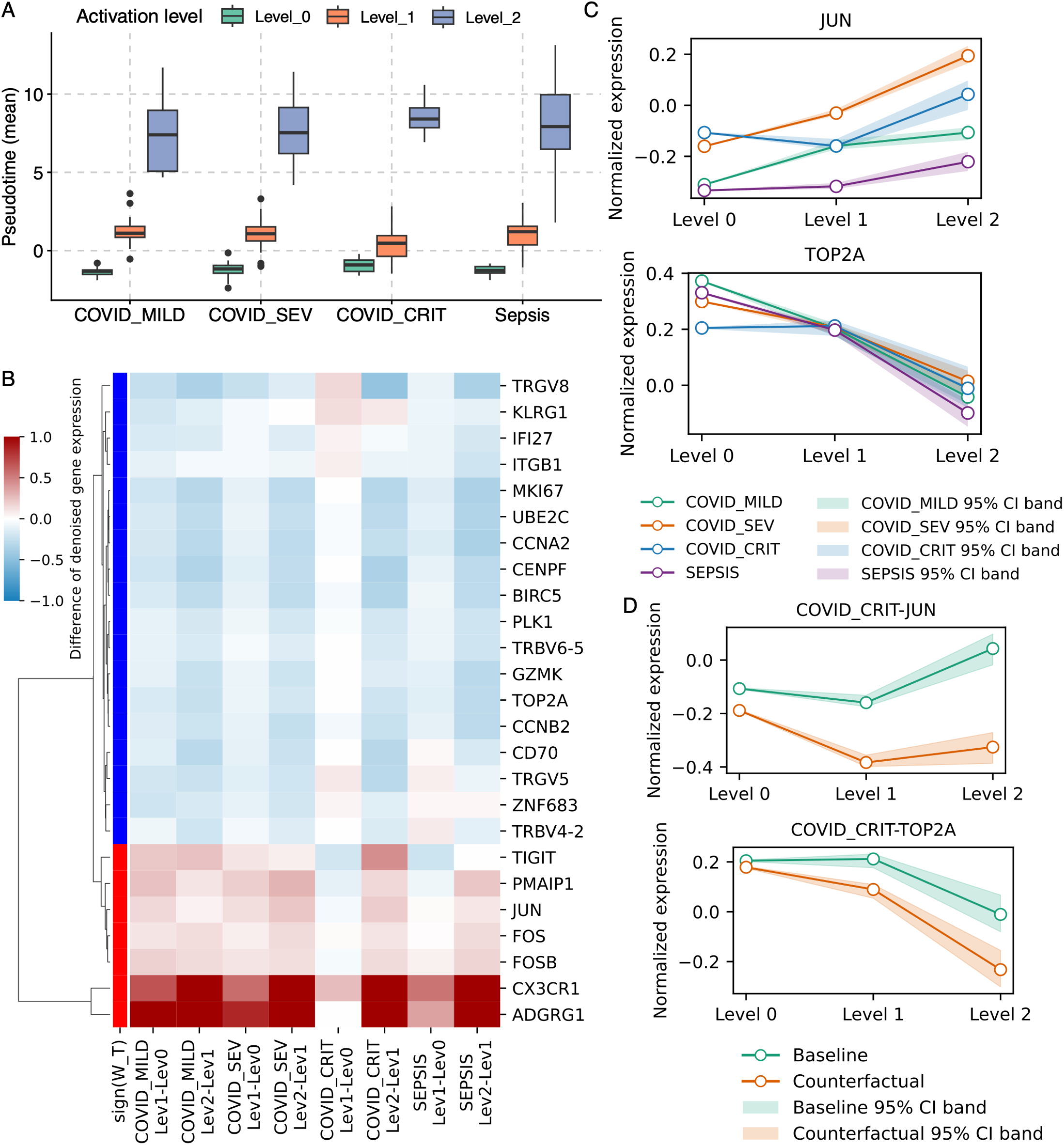
Pseudotime estimation in COVID-19 single-cell RNA data. (A) Posterior median of donor-specific pseudotime coordinates in three CD4 transcriptomic states for COVID-19 severity categories and sepsis. Pseudotime estimates are summarized by the median and interquartile range for each transcriptional state. (B) Differences of pseudotime-associated genes based on 95% credible intervals (CrIs) across transcriptional state transitions with corresponding signs of pseudotime coefficients (*W*_*T*_) that link between gene expression and pseudotime. (C) Estimated denoised gene expression across CD4 transcriptomic states in COVID-19 severity categories and sepsis. Shaded regions represent 95% CrIs. (D) The left panel shows counterfactual denoised gene expression under a counterfactual intervention in which the non-negative matrix factorization (NMF) module 1 of critical COVID-19 samples was increased to the mean level of NMF module 1 observed in mild COVID-19 samples. The right panel shows the difference between counterfactual- and baseline-denoised gene expression. Shaded regions represent 95% CrIs. The red dashed line represents a difference of 0.

To assess the variability of pseudotime intervals across COVID-19 severity and sepsis, we quantified the dispersion of the pseudotime interval between level 0 and level 1. The standard deviations of the intervals were 1.068 (critical COVID-19), 0.858 (severe COVID-19), 0.806 (mild COVID-19), and 1.174 (sepsis). Dispersion differed significantly across COVID-19 severity and sepsis, as assessed by the Brown–Forsythe test (p-value = 0.0036) (**Figure S7D**).

To investigate molecular programs associated with the shortened pseudotime interval, we identified credible genes that were associated with narrowed pseudotime between level 0 and level 1 in critical COVID-19 samples based on 95% CrIs. Using the same criteria applied to the thymus case study, 113 of 500 genes were credibly associated with a narrowed pseudotime window in critical COVID-19 relative to mild COVID-19 and severe COVID-19 groups. These genes included cell-cycle and proliferation-associated genes (e.g., TOP2A, UBE2C), CD4 activation and immediate-early response genes (e.g., JUN, FOS), T cell receptor-related genes (e.g., TRBV4-2, TRBV6-5) (**Figure 4B**). Notably, JUN and FOS showed credibly higher expression at the level 0 state in COVID-critical compared with other groups, with differences credible based on 95% CrIs (**Figure 4C** and **Figure S8**). These genes exhibited stable trajectories across the level 0 to 1 transition in critical COVID-19, whereas they showed increasing trends during the same state transition window in non-critical groups (group differences credible based on 95% CrIs).

In contrast, the proliferation marker TOP2A was lower at level 0 in critical COVID-19 but remained relatively stable during the level 0 to level 1 transition, while other severity groups showed higher initial expression of TOP2A followed by a decreasing trend. These distinct patterns were credible based on 95% CrIs.

### Innate microenvironment context and counterfactual analysis

To characterize the microenvironment, we modeled expression of key innate immune–derived ligands (including IFNG, CCL2–5, CXCL9–10, IL1B) together with five NMF modules summarizing innate immune states. Ranked gene sets based on NMF were used to perform GSEA to annotate each representative immune program (**Table S4**). The top contributing genes for each module are provided in **Figure S9A**. Briefly, NMF0 represents a highly activated myeloid program (e.g., S100A8, S100A9, and LYZ). NMF1 represents proliferating, activated innate-immune and antigen-presenting cell (APC) program (e.g., S100A8, UBE2C, and CD14). NMF2 represents an innate effector program (e.g., GNLY and S100A8). NMF3 represents interferon (IFN) and APC program (e.g., IFITM1 and HLA-DRA). Lastly, NMF4 represents the IFN program (e.g., IFI27 and IFITM1).

Critical COVID-19 samples exhibited elevated NMF0 alongside reduced NMF1, NMF2, and NMF3, consistent with an activated innate inflammatory state coupled with impaired antigen presentation capacity (**Figure S9B**). Estimated associations between pseudotime-associated genes and microenvironmental features are provided in **Figure S9C** as part of the BIOCURRENT framework.

To assess whether these microenvironmental features contribute to altered CD4 pseudotime geometry, we performed counterfactual simulations. Counterfactual reduction of NMF0 in critical COVID-19 was causally associated with a decrease in ΔΔ*T* (-4.247 [95%CrI: -4.248, -4.247]) for the level 0 to level 1 transition between critical COVID-19 and mild COVID-19, compared with baseline. Similarly, counterfactual increase in NMF1 and NMF2 were associated with decreases in ΔΔ*T* of -0.385 [95%CrI: -0.385, -0.385] and -0.169 [95%CrI: -0.169, -0.169], respectively, compared with baseline.

At the gene level, microenvironmental counterfactual interventions showed distinct mechanistic readouts. Counterfactual reduction of NMF0 increased JUN expression in critical COVID-19 at level 0, whereas counterfactual increase of NMF1 or NMF2 reduced JUN in critical COVID-19 at level 0, shifting it toward levels observed in mild COVID-19 cases (**Figure 4D** and **Figure S10**). The same counterfactual interventions also reduced TOP2A across level 0 to level 2 (**Figure S11**).

Together, these results indicate that excessive inflammatory innate activation (NMF0), rather than coordinated antigen-presenting or effector programs (NMF1–2), causally contributes to aberrant CD4 pseudotime geometry in critical COVID-19 phenotype. Counterfactual attenuation of this inflammatory microenvironment partially normalized pseudotime transitions between naive and effector CD4 states, accompanied by gene-level shifts in transcriptional regulation and cell-cycle activity.

## DISCUSSION

Numerous pseudotime inference methods have been developed to infer transcriptomic coordinates in scRNA and spatial transcriptomic platforms^8^. BIOCURRENT extends the transcriptomic coordinate inference by incorporating available baseline characteristics and microenvironment to alleviate model mis-specification in pseudotime identification. BIOCURRENT provides not only generative pseudotime coordinates from scRNA-seq or spatial transcriptomic datasets, but also enables counterfactual simulations to test whether microenvironmental perturbations lead to intracellular transcriptomic alterations and directional changes in pseudotime geometry.

Advanced transcriptomic profiling provides an unprecedented opportunity to uncover disease pathogenesis at cellular resolution. However, individuals exhibit substantial heterogeneity not only in baseline characteristics but also in their microenvironmental contexts. In such settings, population-based comparisons may obscure core biological alterations relevant to therapeutic translation^27,28^. In this study, we demonstrated that distinct patient groups are characterized by compressed or extended pseudotime coordinates between specific adjacent transcriptional states, such as naive-to-effector or DP early to DP late transitions. These differences define state-specific alterations, indicating where the transcriptional landscape becomes constrained or prematurely primed. Importantly, such localization of pseudotime deviation informs therapeutic windows by identifying whether shifts in transcriptomic programs emerge early or later in the transcriptomic coordinates and whether upstream or preventive intervention may be feasible. These trajectory-geometry features reflect early transcriptional readiness, inflammatory priming, and microenvironmental skewing, and these features are not detectable through differential gene expression analysis alone.

BIOCURRENT was designed to formalize and investigate the pseudotime heterogeneity in pseudotime geometry. By jointly modeling baseline covariates and microenvironmental context, BIOCURRENT treats trajectory geometry itself as a quantitative phenotype. In COVID-19 scRNA-seq data, the sepsis group exhibited greater variability in pseudotime intervals between CD4 naive to effector states, compared with COVID-19 severity categories. These findings suggest that, in future analyses, patients may be stratified according to geometry-derived features to evaluate whether such patterns are associated with downstream clinical trajectories, including vulnerability or resilience to disease progression.

In conclusion, we introduce BIOCURRENT, a causal inference framework for advanced transcriptomics that infers pseudotime geometry and identifies individual- and condition-specific alterations in pseudotime coordinates. By modeling gene regulation as a function of baseline covariates, microenvironmental context, and latent pseudotime, BIOCURRENT supports counterfactual simulation of upstream microenvironmental modulation and quantifies its impact on state-specific trajectory geometry through the ΔΔ*T* estimator. Applications to thymic development and COVID-19 immune dysregulation demonstrate that heterogeneity in pseudotime intervals localizes state-specific trajectory alterations, and that these alterations can be partially alleviated through counterfactual modulation of microenvironmental niches. We anticipate that BIOCURRENT will provide a valuable foundation for investigating state-aware transcriptional phenotypes, enabling mechanistic hypothesis generation to guide future experimental and clinical studies.

### Limitations of the study

BIOCURRENT models gene expression as a linear function of latent pseudotime. This design prioritizes causal identifiability and interpretability of the latent pseudotime coordinate over maximal predictive flexibility. While non-linear pseudotime–to-gene expression relationships may better fit complex transcriptional patterns, such flexibility can render the latent pseudotime non-identifiable.

Second, BIOCURRENT estimates pseudotime as a latent variable inferred from observed gene expression. Because pseudotime is a fully latent coordinate without an external temporal anchor, counterfactual pseudotime under modulation of upstream contextual variables is not directly defined, and re-timing of the latent trajectory is therefore not identifiable. To address this conceptual limitation, BIOCURRENT introduces the ΔΔ*T* estimator, which quantifies the direction and magnitude of context-dependent distortion in state transition intervals along the inferred pseudotime geometry, providing an interpretable surrogate for relative acceleration or compression of biological progression without requiring counterfactual re-timing of the latent coordinate itself.

Third, BIOCURRENT does not explicitly model gene–gene regulatory interactions. Genes are treated as conditionally independent given baseline characteristics, latent pseudotime, and microenvironmental features. While explicit modeling of co-regulatory modules or pathway structures may better reflect biological co-regulation, such approaches require assuming that regulatory groupings remain invariant under counterfactual intervention. Because regulatory modules themselves may reorganize across conditions or microenvironmental perturbations, imposing fixed co-regulatory clusters risks introducing bias into counterfactual estimation.

Finally, BIOCURRENT focuses on estimating how progression geometry and state transition intervals vary across conditions or donors, rather than inferring lineage topology. This design aligns with immune and developmental settings in which the central scientific question concerns differential progression along a dominant biological program, rather than fate choice itself. In addition, explicit inference of branching structure can introduce substantial uncertainty and non-identifiability of latent progression coordinates, particularly in cross-sectional transcriptomic data. Future extensions of BIOCURRENT may incorporate branching-aware pseudotime representations or multi-trajectory coupling, enabling unified analysis of trajectory topology and condition-specific progression geometry within a single probabilistic framework.

## Materials availability

This study did not generate new materials.

## Data and code availability

- The datasets that are analyzed within the current study are publicly available.
- BIOCURRENT is publicly available as a Python package on Github (https://github.com/Kamaleswaran-Lab/biocurrent). Notebooks for running all of the analysis performed in this manuscript are available.
- Any additional information required to reanalyze the data reported in this paper is available from the lead contact upon request.

## ACKNOWLEDGMENTS

The authors thank Min Huang, Chengkun Yan, Yifang Xi for their helpful discussion. S.K. was supported by the National Institutes of Health (NIH) under Award Number R21GM148931 and R01HL170175. C.M.C was supported by NIH under Award Number R35GM148217. R.K. was supported by the NIH under Award Numbers R01GM139967, R21GM151703, R21GM148931, and R01HL170175.

## AUTHOR CONTRIBUTIONS

S.K., S.A.R., S.P.R, and R.K. conceptualized the research direction. S.K. and R.K. developed methodology and performed analyses. S.K. and R.K. wrote the original manuscript and S.A.R., S.P.R, C.M.C and R.K. reviewed the manuscript.

## SUPPLEMENTAL INFORMATION INDEX

Table S1-S2 and Figures S1-S11 and their legends in a PDF

Table S3. Gene set enrichment analysis of microenvironmental modules derived from non-negative matrix factorization in thymic early T-cell developmental progression in an Excel file

Table S4. Gene set enrichment analysis of microenvironmental modules derived from non-negative matrix factorization in CD4 transcriptomic states in COVID-19 in an Excel file

## METHODS

### BIOCURRENT

BIOCURRENT models observed gene expression as a function of a latent transcriptomic coordinate (pseudotime), without assuming a strict ordering of individual observations. Let *G*_*g,i*_ denote the observed expression of gene *g* in observation *i* (e.g., a cell or a bulk RNA-seq sample). Each observation *i* is associated with a latent pseudotime coordinate *T*_*i*_, where multiple observations may share identical or similar *T*_*i*_ values. Baseline covariates are denoted by *Z*_*i*_ (e.g., age, sex), microenvironmental features by *L*_*i*_ (e.g., ligand expression or tissue architectural variables), and sequence-related technical covariates by *Q*_*i*_.

We model gene expression using the following linear formulation:

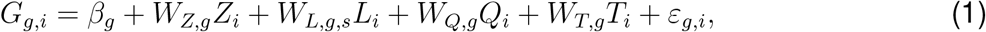

where *β*_*g*_ is a gene-specific intercept. *W*_*Z,g*_ ∈ ℝ^*z*^ denotes gene-specific coefficients for baseline covariates, where *z* is the number of baseline covariate nodes in *Z. W*_*L,g,s*_ ∈ ℝ^*l*^ denotes gene (*g*)- and transcriptional state (*s*)-specific coefficients for microenvironmental features, where *l* is the number of microenvironmental nodes in *L. W*_*Q,g*_ ∈ ℝ^*q*^ denotes gene-specific coefficients for technical covariates, where *q* is the number of nodes in *Q. W*_*T,g*_ ∈ ℝ captures the sensitivity of gene *g* to variation along the latent pseudotime coordinate. The residual term *ε*_*g,i*_ represents observation-specific exogenous noise within the structural causal model^29^.

### Parameter priors

The structural parameters, including *W*_*Z*_, *W*_*Q*_, and *W*_*T*_, were defined using Gaussian priors:

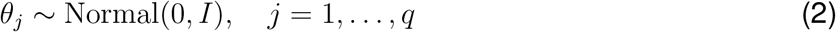

where *θ* = (*θ*_1_, …, *θ*_*q*_) collects the structural parameters.

For the parameter *W*_*L,s*_, transcriptional state-specific coefficients that link microenvironmental context to gene expression, we place Laplace priors to encourage sparsity:

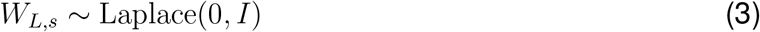

Exogenous noise terms (*ε*_*g,i*_), representing observation-level gene expression variability, were modeled using low-rank latent factors with Gaussian priors. This formulation introduces structured correlation across genes and cells in the residual variation. It is both biologically realistic and computationally efficient, given a large number of observations (cells) in the observed datasets.

### Pseudotime constraints

BIOCURRENT incorporates weak structural constraints to improve the identifiability of transcriptomic geometry. Specifically, users provide a set of ordered transcriptional states, such as naive → effector → terminal or double-negative → double-positive states. For each donor–state combination, BIOCURRENT assigns a state-specific pseudotime mean parameter, with these means constrained to follow the specified order within each donor. Thus, different donors may have different pseudotime means for the same transcriptional state. Within a given donor–state, observations (cells) share the same state-level pseudotime mean, while allowing cell-level variability around this mean. This design anchors the latent pseudotime axis to biologically interpretable states within individuals, without enforcing strict cell-level ordering or cross-donor alignment, thereby allowing inter-individual shifts in transcriptomic coordinates. Priors of pseudotime parameters are centered at zero; therefore, these constraints do not enforce differences across donors or transcriptional states.

### Learning scheme

BUICURRENT learns latent transcriptomic coordinates using stochastic variational inference, implemented in Pyro^30^, a probabilistic programming framework built on PyTorch^31^. To mitigate the risk of latent variable collapse, we applied Kullback–Leibler (KL) annealing^32^, a training strategy that gradually introduces the regularization term during optimization. Early training emphasizes fitting the observed data, while the strength of the regularization is progressively increased to encourage stable latent representations. In our implementation, the annealing weight was increased from 0.001 to 1.0 over the first 50% of training iterations following a modified exponential schedule.

Optimization was performed using the Adam optimizer with the `ReduceLROnPlateau` learning-late scheduler in PyTorch^31^. The initial learning rate of 0.03 was reduced by a factor of 0.9 when the loss did not improve for 100 consecutive iterations.

### Adjustment for unbalanced donor and transcriptional states

In single-cell RNA-seq data, the number of recovered cells often varies across donors, and the proportions of specific transcriptional states may differ between conditions. This imbalance is particularly pronounced in T cell subsets in sepsis, where lymphopenia leads to severe under- representation of certain states compared with non-sepsis donors.

To prevent the majority class from dominating the parameter learning, BIOCURRENT incorporates per-donor and per-transcriptional-state weighting in the learning objective. Specifically, each donor–state combination is assigned a weight proportional to the inverse of its frequency in the observed data. These weights are applied to the likelihood term during optimization, ensuring that observations from under-represented donors or transcriptional states contribute to parameter estimation.

### Two-stage learning for parameters

BIOCURRENT distinguishes between structural parameters governing gene regulation and the exogenous noise terms in the model. To prevent the noise component from absorbing the structured signal, we adopt a two-stage learning strategy inspired by causal modeling principles^33^.

In the first stage, structural parameters are optimized while the exogenous noise is held fixed. This encourages the model to explain systematic variation in gene expression through baseline covariates, microenvironmental features, and latent pseudotime, rather than through noise. In the second stage, the learned structural parameters are held fixed, and the noise model is relaxed, allowing the exogenous noise terms to capture residual variability not explained by the structural components.

For each stage, optimization was performed using mini-batch stochastic variational inference. In the first stage, 2,000 observations (cells) were randomly sampled per iteration for 100 epochs. In the second stage, 1,000 observations were sampled per iteration for 10 epochs.

### Synthetic data experiments

To evaluate the identifiability of donor-specific pseudotime coordinates and associated model parameters, we simulated multi-donor single-cell RNA-seq data using the BIOCURRENT generative structure, with predefined ground-truth parameters.

We simulated data for five donors, each comprising *M* ordered transcriptional states. Each transcriptional state contained *N* cells and 500 genes. Microenvironmental covariates consisted of 20 features. We varied the levels of sparsity of the coefficients that link microenvironmental context to gene expression to reflect heterogeneous and partially irrelevant environmental effects. Exogenous noise was added to each gene expression for each cell.

We considered the following simulation settings:

- Number of ordered transcriptional states: *M* ∈ *{*4, 8*}*;
- Number of cells per transcriptional state: *N* ∈ *{*100, 500*}*;
- Sparsity of microenvironmental coefficients: 95% and 99%.

For each combination of *M*, *N*, and sparsity level, we generated 50 independent replicate datasets.

### Performance metrics in synthetic data experiments

Parameter recovery for denoised gene expression was evaluated using root mean squared error (RMSE) between true and estimated values, divided by the standard deviation of the true parameter distribution (normalized RMSE). Pseudotime recovery was assessed using Pearson’s correlation coefficient between true and inferred pseudotime coordinates, which was evaluated both globally across all donors and separately at the donor level. These simulations explicitly test BIOCURRENT’s ability to disentangle donor-specific shifts in pseudotime geometry from microenvironmental and stochastic variation.

### Benchmarking BIOCURRENT using time-stamped transcriptomic data

To evaluate the fidelity of BIOCURRENT in recovering true temporal coordinates from transcriptomic data, we analyzed a publicly available time-series microarray dataset (GSE39598), which profiles gene expression in a single *Arabidopsis thaliana* leaf following *Botrytis cinerea* infection^21^. Gene expression was measured every two hours up to 48 hours post-infection, yielding 24 time-stamped samples.

To align with the prior pseudotime inference setting^14^, we grouped the 24 samples into four ordered transcriptomic states, each containing six consecutive time points. Raw expression values were pseudo-log normalized and standardized for each feature, and 150 highly variable genes were selected using Scanpy^34^.

Using the four ordered transcriptomic states as input, BIOCURRENT was applied to infer latent pseudotime coordinates, which were then compared against the true sampling times. For benchmarking, we evaluated BIOCURRENT alongside representative pseudotime inference methods, including nearest-neighbor–based, diffusion-based, and tree-based approaches. Specifically, Slingshot^13^, Diffusion Pseudotime (DPT)^9^, and TSCAN^12^.

### Real-world data application: thymus spatial transcriptomics

To demonstrate BIOCURRENT application, we downloaded publicly available healthy fetal and postnatal thymus spatial transcriptomic datasets^18^ from the CellxGene data portal^35^. The original study provided multiple biological replicates. To avoid duplicated samples and to ensure sufficient spatial coverage, we retained unique donors, resulting in eight fetal samples and four postnatal samples.

Thymus spatial transcriptomic data were preprocessed. Specifically, we filtered spots to those with 200-8,000 genes and excluded spots with a mitochondrial gene proportion more than 15%. Spatial deconvolution was then performed using cell2location^23^ to estimate cell-type-specific gene expression profiles.

As a reference for spatial deconvolution, we used preprocessed healthy thymus scRNA-seq data (GSE271304)^18^ and downloaded from the same CellxGene collection^35^. The granularity of the deconvolution depends on the reference cell type annotation. The original thymus scRNA-seq data provided four levels of a hierarchical annotation scheme, primarily organized by transcriptional cell states rather than classical thymocyte developmental stages (e.g., DN1, DN2). The most granular annotation level, cell type level 4 explore, included progenitor T cells as well as early and late phases of double-positive (DP) thymocytes. We therefore used this most granular annotation level as the reference and subsequently aggregated cell types (**Table S1**) to align them with canonical T cell developmental trajectories. Following spatial deconvolution, we constructed a pseudo-single-cell dataset by aggregating the deconvoluted cell type-specific gene expression estimates across spatial spots. Pseudo-cells with fewer than 1,500 expressed genes, defined as genes with expression values above the median expression across spatial spots, were excluded, resulting in 17,365 pseudo-cells. Principal component analysis (PCA) of the gene expression matrix revealed no technical batch effect. To assess the biological plausibility of cell2location-derived cell type–specific gene expression estimates, we evaluated the expression of canonical marker genes using the pseudo–single-cell dataset, with confirmation that cell type-specific gene expression profiles reasonably represent cortical T cell developmental lineages for our BIOCURRENT application.

### Preprocessing of thymus spatial transcriptomic data

For the first real-world data application of BIOCURRENT, we inferred pseudotime coordinates across T cell developmental lineages, including early T cell progenitors (ETP), double-negative (DN) early, DN late, double-positive (DP) early, and DP late stages. We retained only spatial spots annotated as cortical regions in the original study. Mitochondrial genes, ribosomal genes, hemoglobin-associated genes, and genes without an assigned gene symbol were excluded. Subsequently, 500 highly variable genes across all samples were selected.

To represent the microenvironment, we extracted cortical thymic epithelial cell (cTEC)–specific gene expression profiles for each spatial spot and selected 1,000 highly variable genes. Among these genes, key cTEC-derived ligands (DLL4, CXCL12, CCL25, and WNT4) were retained as individual features, while the remaining genes were summarized into five modules using non-negative matrix factorization (NMF) with scikit-learn^36^.

Baseline covariates included biological sex and developmental age, expressed as gestational weeks centered at 40 weeks. Sequence-related technical covariates included `log1p_total_counts_log1p_n_genes_by_counts`, and `percent_mito`. Single-cell RNA-seq data preprocessing was performed using the Python package Scanpy^34^.

### Real-world data application: COVID-19 scRNA-seq data

For the second real-world data application of BIOCURRENT, we analyzed publicly available single-cell RNA-seq data from the **CO**vid-19 **M**ulti-omics **B**lood **AT**las (COMBAT) consortium^2^, downloaded via the CellxGene platform^35^. To investigate pseudotime geometry across disease severity, we included samples from mild COVID-19, severe COVID-19, critical COVID-19, and sepsis samples, with sepsis defined as all-cause sepsis excluding SARS-CoV-2 infection. COVID-19 severity was defined according to World Health Organization (WHO) guidelines and annotated by the original study. For donors with multiple blood samples, only the earliest available sample was retained. After normalization and scaling, PCA was performed to assess potential technical batch effects, and no substantial batch effects were observed.

We modeled CD4 T cell activation along three ordered transcriptional stages: level 0 (naive), level 1 (effector), and level 2 (terminal), following the original cell-type annotations (**Table S2**). Enrichment of canonical CD4 T cell markers, including CD3D and CD4, was confirmed for each stage (**Figure S7A**). Mitochondrial genes, ribosomal genes, hemoglobin-associated genes, and genes without assigned gene symbols were excluded. Subsequently, 500 highly variable genes across all samples were selected.

To represent microenvironmental features reflecting circulating innate immune context, we extracted gene expression profiles from classical monocytes, non-classical monocytes, dendritic cells, and natural killer cells (**Figure S7B**). Key ligands and cytokines involved in innate–adaptive immune interactions were retained as individual features, including IFNG, IL1B, CXCL9, CXCL10, CCL2, CCL3, CCL4, and CCL5. The remaining 992 genes among the 1,000 most variable innate-associated genes were summarized into five modules using non-negative matrix factorization (NMF).

Baseline covariates included age, sex, ethnicity, pre-existing comorbidities, smoking history, and blood collection timing relative to symptom onset, encoded as 19 dummy variables. To improve the identifiability of pseudotime and maintain orthogonality among covariates, baseline features were projected using PCA. We retained the first three PCs as determined by scree plot inspection.

Sequence-related covariates included `QC_ngenes`, `QC_total_UMI`, `QC_pct_mitochondrial`, and `QC_scrub_doublet_scores`.

### Ethical Statement

The datasets used in this study, including *Arabidopsis thaliana* leaf bulk RNA-seq, thymus spatial transcriptomics, and COMBAT scRNA-seq, are publicly available and de-identified. For the human-derived datasets (thymus and COMBAT), the use of these datasets was reviewed by the Emory Institutional Review Board and determined to qualify as non-human subjects research. All study procedures were conducted in accordance with the ethical principles of the Declaration of Helsinki.

### Quantification and statistical analysis

#### Pseudotime-associated gene identification and definition of ΔΔ*T*

In our BIOCURRENT formulation, the gene-specific pseudotime coefficient *W*_*T,g*_ (for gene *g*) represents the association between gene expression and pseudotime after accounting for sequence-related covariates, baseline characteristics, and microenvironmental context. Therefore, for genes credibly associated with pseudotime coordinates, the 95% credible intervals of *W*_*T,g*_ should exclude zero. Following the BIOCURRENT formulation, changes in denoised gene expression across transcriptional states can be interpreted through the pseudotime axis defined by *W*_*T,g*_. For a given gene *g*, let 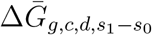 be the change in mean denoised gene expression between two transcriptional states *s*_1_ and *s*_0_ given a condition *c* and donor *d*. We define a gene-level estimator of pseudotime interval as:

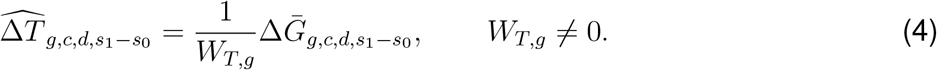

This estimator can be interpreted as a projection of the denoised gene-expression change onto the pseudotime axis defined by *W*_*T,g*_, rather than as an exact decomposition isolating pseudotime-specific effects from microenvironmental contributions.

Subsequently, the contrast of 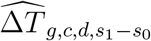 across conditions *c*_1_ and *c*_0_ is defined as:

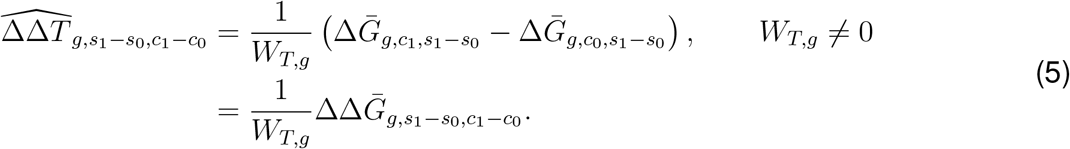

When the contrast of gene-expression changes between transcriptional states across conditions 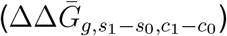 is aligned with pseudotime variation, its sign is expected to be concordant with the sign of the gene-specific pseudotime coefficient *W*_*T,g*_. Conversely, sign discordance indicates that other factors, such as microenvironmental or baseline characteristic contributions, are sufficiently large to offset the association between pseudotime and gene expression.

Accordingly, a gene *g* was considered contributory to a pseudotime interval if it satisfied the following criteria:

1. 0 ∈*/* CrI_95%_(|*W*_*T,g*_|);

2. 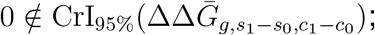

3.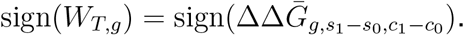.

Using Eq. 5, we define the aggregated ΔΔ*T* estimator over credible genes associated with pseudotime as:

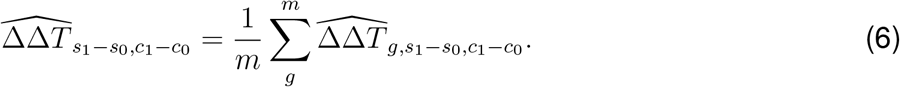

where *m* is the number of credible genes associated with pseudotime coordinates.

In counterfactual analysis, microenvironmental programs are perturbed to obtain counterfactual gene-expression estimates. The counterfactual effect is defined as the change in the ΔΔ*T* estimator relative to baseline:

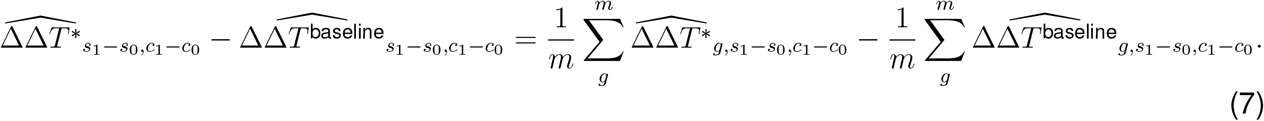

where ^∗^ denotes counterfactual quantities.

#### Statistical test for differences of pseudotime intervals

To compare pseudotime interval differences across conditions, we performed statistical tests on donor-level interval summaries, treating donors as independent units, specifically, the posterior median of pseudotime for each donor. Because these summaries were not assumed to follow a normal distribution, we used Welch’s t-test (for two conditions) or Welch’s one-way ANOVA (for more than two conditions) for interpretability and computational simplicity. Welch’s one-way ANOVA was followed by pairwise comparisons using the Games–Howell post hoc test to control the family-wise error rate across all contrasts. The significance threshold was set at *α* = 0.05.

