## Supplemental material for "Microenvironment-informed inference of transcriptional progression geometry"

Supplementary Table S1: Early T cell developmental state definition based on the original cell types

| **T cell developmental state** | **Original cell type annotation (cell_type_level_4_explore)** |
|---|---|
| T_ETP | T_ETP |
| DN early | T_DN(early), T_DN(P) |
| DN late | T_DN(Q)-intermediate, T_DN(Q) |
| DP early | T_DP(P), T_DP(Q)-early |
| DP late | T_DP(Q), T_DP(Q)-late_vdj |

Supplementary Table S2: CD4 T cell transcriptomic state definition based on the original cell types

| **CD4 transcriptional state** | **Original cell type annotation (cluster)** |
|---|---|
| Level 0 | CD4.NAIVE.1, CD4.NAIVE.2, CD4.NAIVE.3 |
| Level 1 | CD4.TEFF.TCF7, CD4.TEM, CD4.Th, CD4.Th1.1, CD4.Th1.2, CD4.Th1.3, CD4.Th17, CD4.Th1/Th17, CD4.TEFF.GZMK, CD4.TEM.GZMK, CD4.Th.CCR4.CCR10, CD4.Th.CXCR5.KLRB1, CD4.Th.mitohi, CD4.TEM.mitohi.1, CD4.TEM.mitohi.2, CD4.Th.IFN.resp, CD4.TEM.IFN.resp |
| Level 2 | CD4.TEMRA, CD4.TEMRA.KLRB1, CD4.TEMRA.XCL2, CD4.TEMRA.KLRC3, CD4.TEMRA.mitohi |

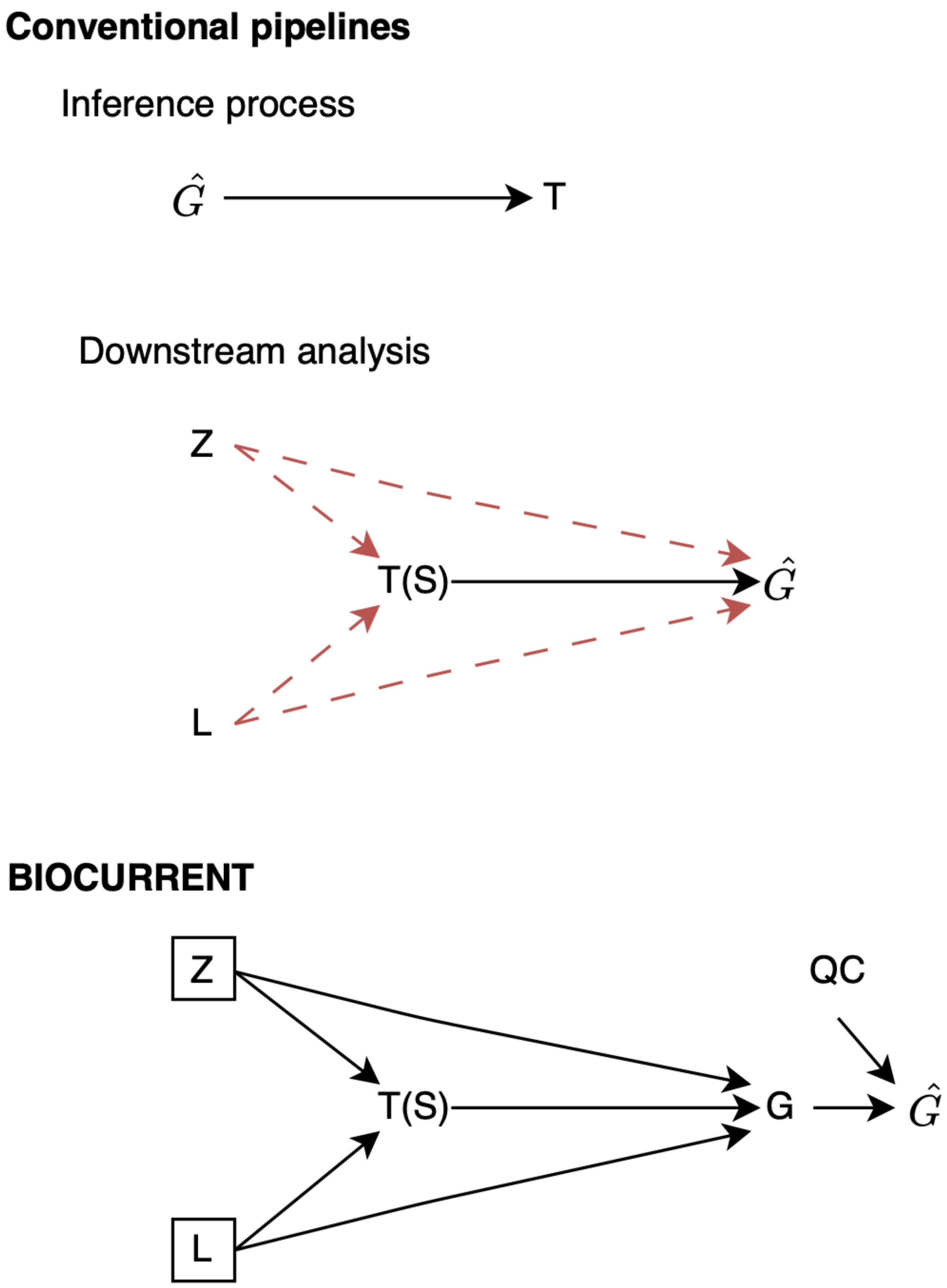


Supplementary Figure S1: The diagram represents conventional pseudotime inference pipelines, specifically, estimating pseudotime $T$ using observed gene expression $\hat{G}$(top) and using the pseudotime for downstream analysis, such as differential gene analysis associated with inferred trajectory or pseudotime (middle). $Z$ denotes baseline covariates and $L$ denotes microenvironmental features. $T(S)$ represents pseudotime $T$ and biological developmental state ($S$) are interchangeably encoded in the downstream analysis. BIOCURRENT constructs latent pseudotime coordinates and estimates denoised gene expression $G$, while conditioning on baseline covariates, microenvironmental covariates, and technical covariates.

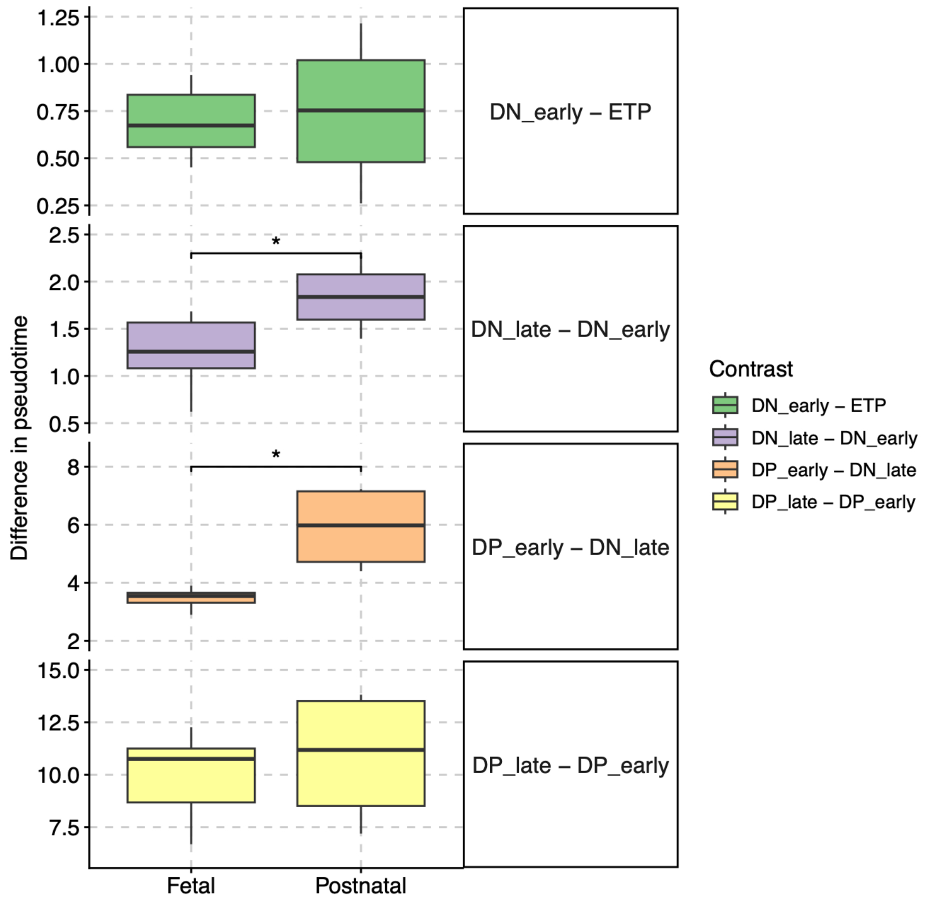


Supplementary Figure S2: Posterior median of donor-specific pseudotime coordinates in five T cell developmental lineages for fetal and postnatal samples. Pseudotime estimates are summarized by the median and interquartile range for each transcriptional state. * denotes statistically significant contrast using Welch's t-test.

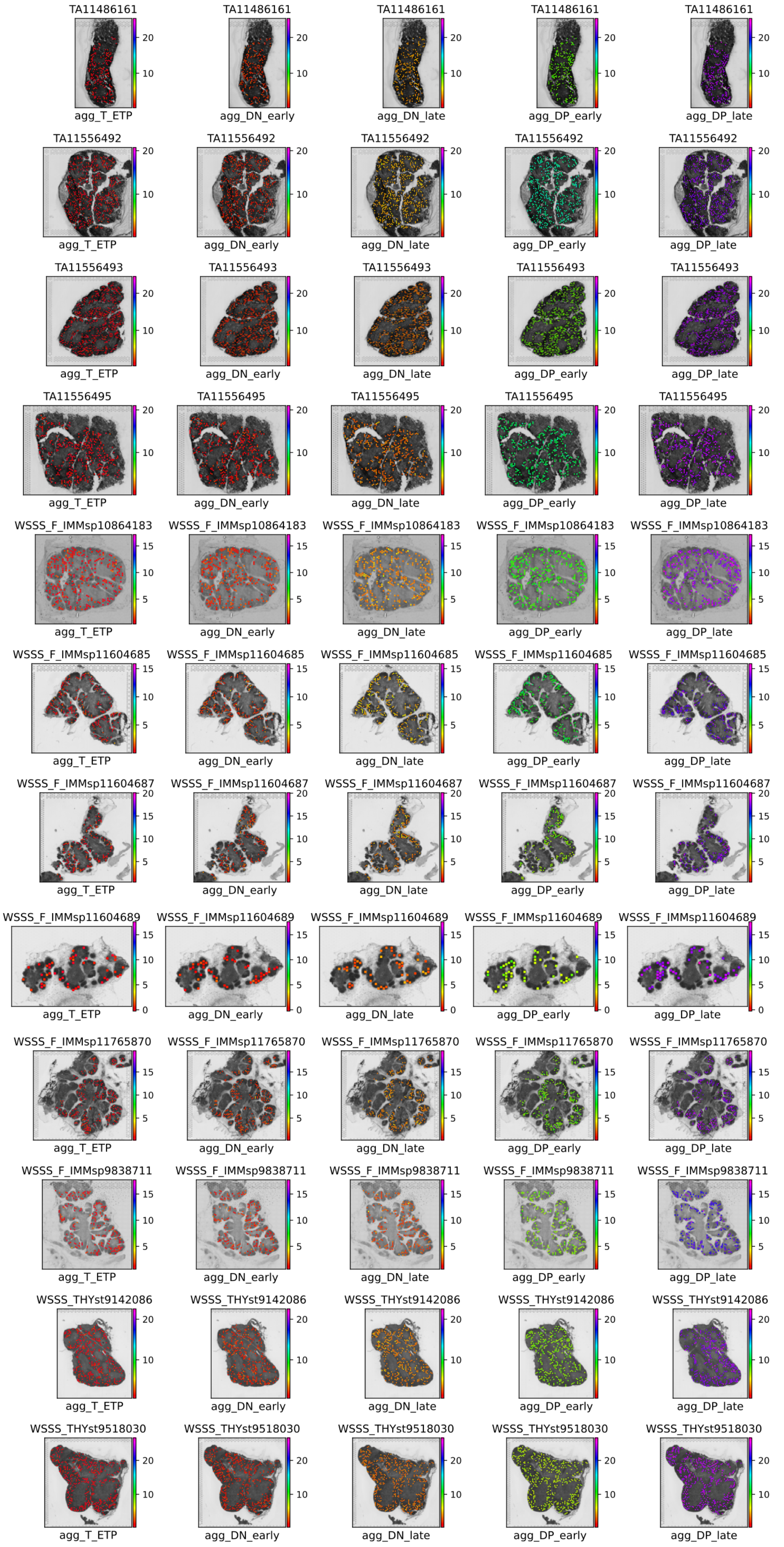


Supplementary Figure S3: Posterior median of pseudotime coordinates were projected onto spatial coordinates of thymic samples. Sample IDs starting with “TA” indicate postnatal samples, whereas “WSSS” indicates fetal samples.

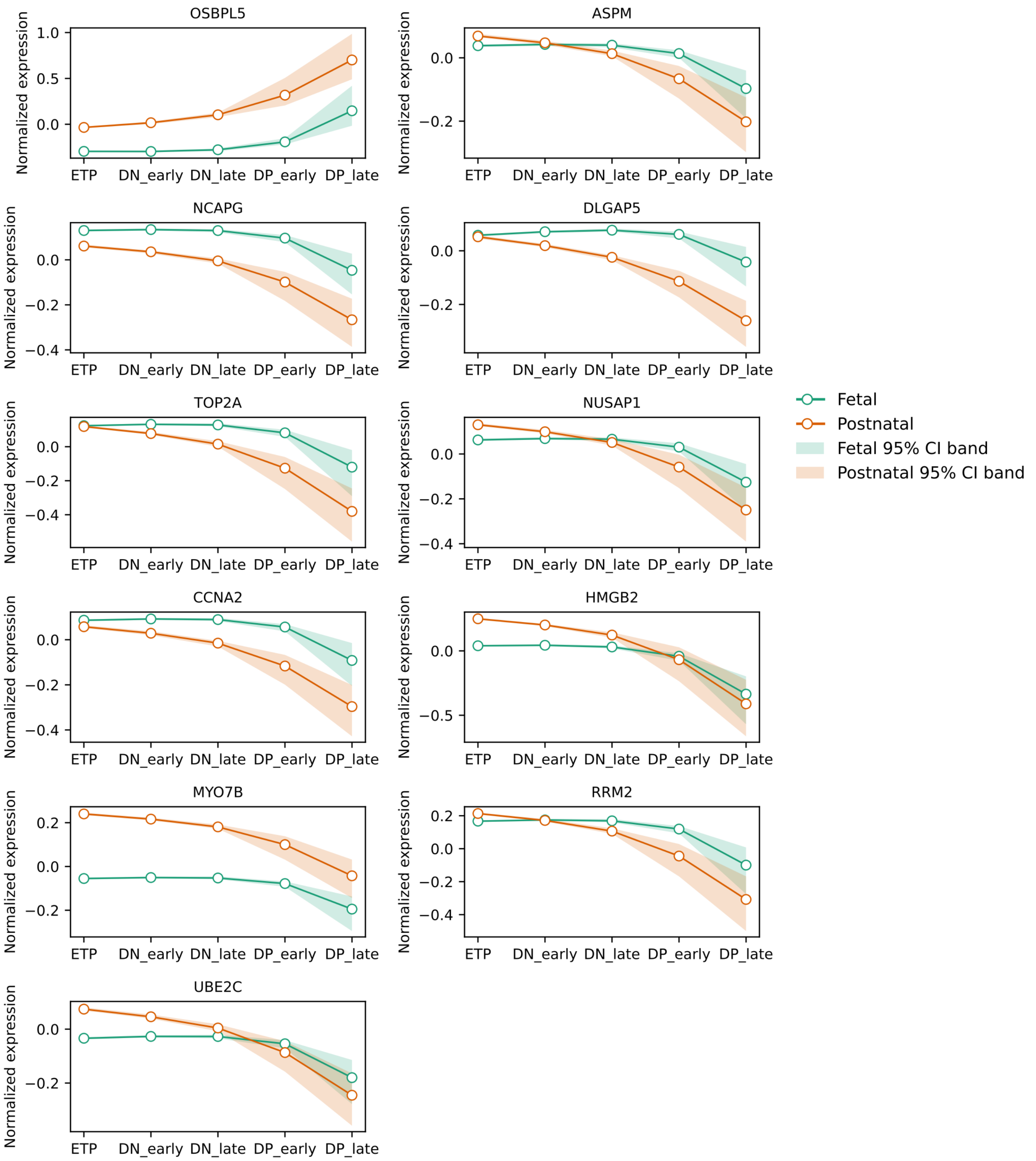


Supplementary Figure S4: Estimated denoised gene expression across T cell developmental lineages for fetal (orange) and postnatal (green) samples. Shaded regions represent 95% credible intervals.

A

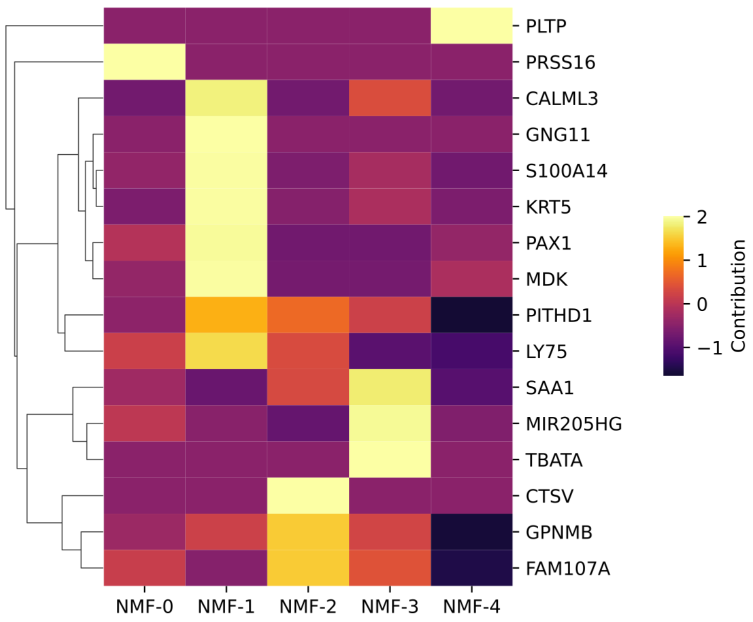

PLTP
PRSS16
CALML3
GNG11
S100A14
KRT5
PAX1
MDK
PITHD1
LY75
SAA1
MIR205HG
TBATA
CTSV
GPNMB
FAM107A
NMF-0
NMF-1
NMF-2
NMF-3
NMF-4
2
1
0
−1
Contribution


B

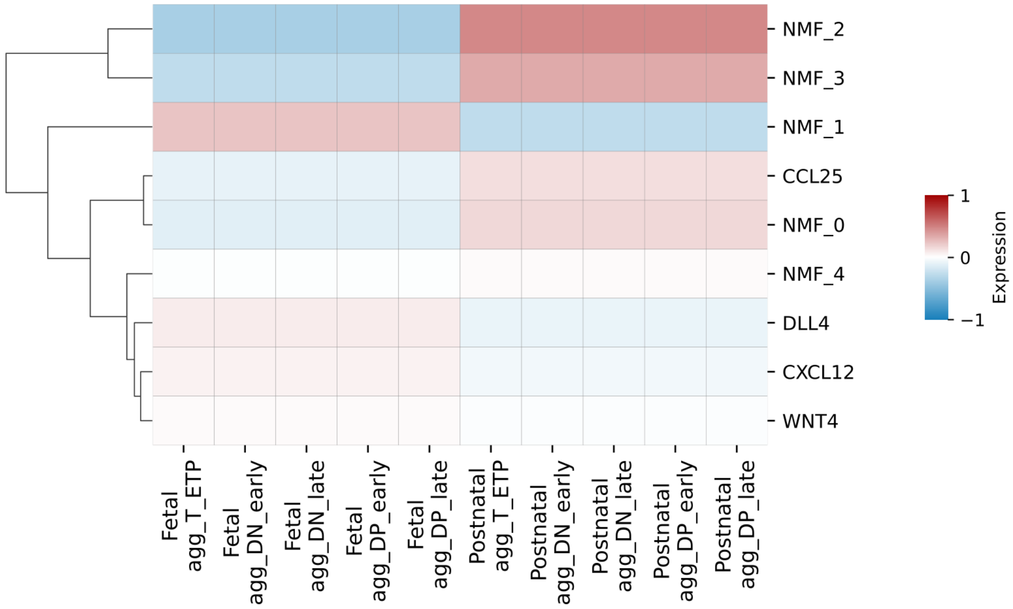

NMF_2
NMF_3
NMF_1
CCL25
NMF_0
NMF_4
DLL4
CXCL12
WNT4
Fetal agg_T_ETP
Fetal agg_DN_early
Fetal agg_DN_late
Fetal agg_DP_early
Fetal agg_DP_late
Postnatal agg_T_ETP
Postnatal agg_DN_early
Postnatal agg_DN_late
Postnatal agg_DP_early
Postnatal agg_DP_late
1
0
−1
Expression


C

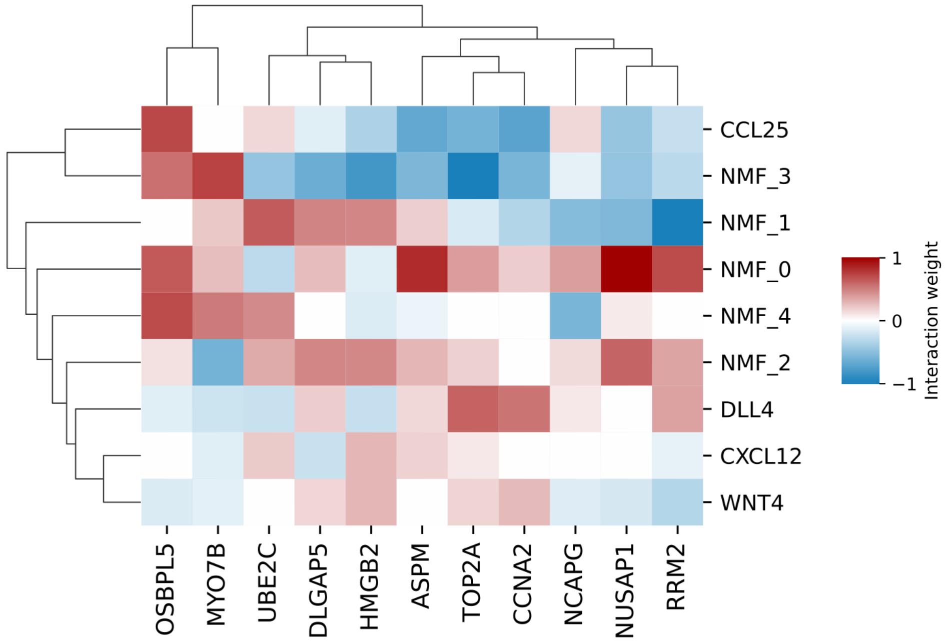

CCL25
NMF_3
NMF_1
NMF_0
NMF_4
NMF_2
DLL4
CXCL12
WNT4
OSBPL5
MYO7B
UBE2C
DLGAP5
HMGB2
ASPM
TOP2A
CCNA2
NCAPG
NUSAP1
RRM2
1
0
−1
Interaction weight

Supplementary Figure S5: Microenvironmental features used to model pseudotime in thymic T cell development. (A) Normalized gene weights (H matrix) for five non-negative matrix factorization (NMF) modules. Yellow indicates relatively higher gene weights within a module, and black indicates lower gene weights after normalization. (B) NMF-derived module activity and gene expression of microenvironmental features across T cell developmental lineages in fetal and postnatal samples. Red indicates a credibly positive change across lineages based on 95% credible intervals (CrIs), and blue indicates a negative change. Features with non-credible changes are shown in white. (C) Estimated microenvironmental coefficients linking microenvironmental features to pseudotime-associated genes. Red indicates a credibly positive association based on 95% CrIs, and blue indicates a negative association. Non-credible pairs are shown in white.

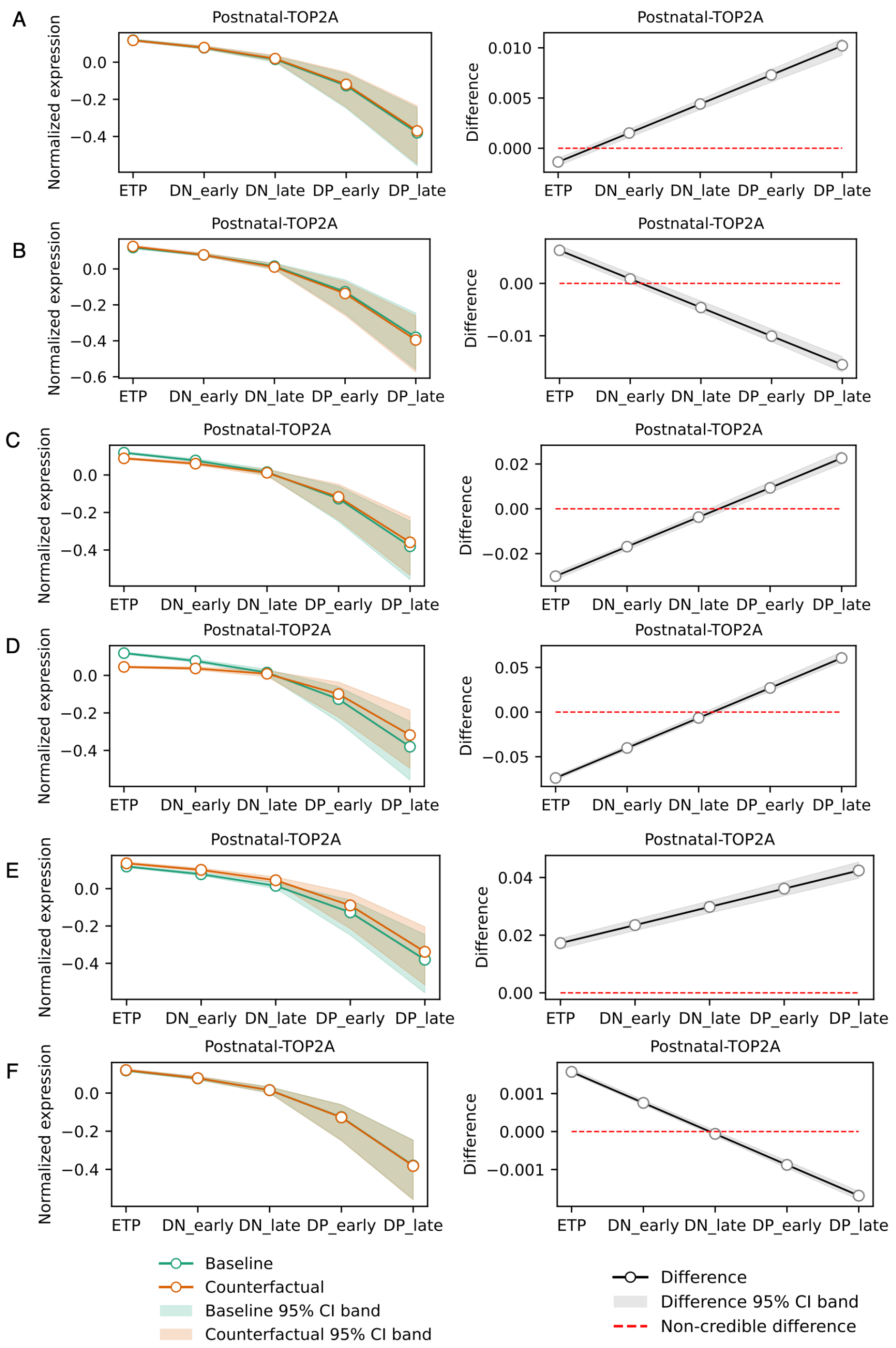


Supplementary Figure S6: Counterfactual intervention on postnatal samples by modulating microenvironmental features to the mean levels observed in fetal samples. Left panels show baseline and counterfactual denoised gene expression, with shaded regions indicating 95% credible intervals (CrIs). Right panels show the difference between counterfactual and baseline gene expression, with shaded regions indicating 95% CrIs. (A) Counterfactual increase of DLL4 to the mean level observed in fetal samples. (B) Counterfactual decrease of NMF0 to the mean level observed in fetal samples. (C) Counterfactual increase of NMF1 to the mean level observed in fetal samples. (D) Counterfactual decrease of NMF2 to the mean level observed in fetal samples. (E) Counterfactual decrease of NMF3 to the mean level observed in fetal samples. (F) Counterfactual decrease of NMF4 to the mean level observed in fetal samples.

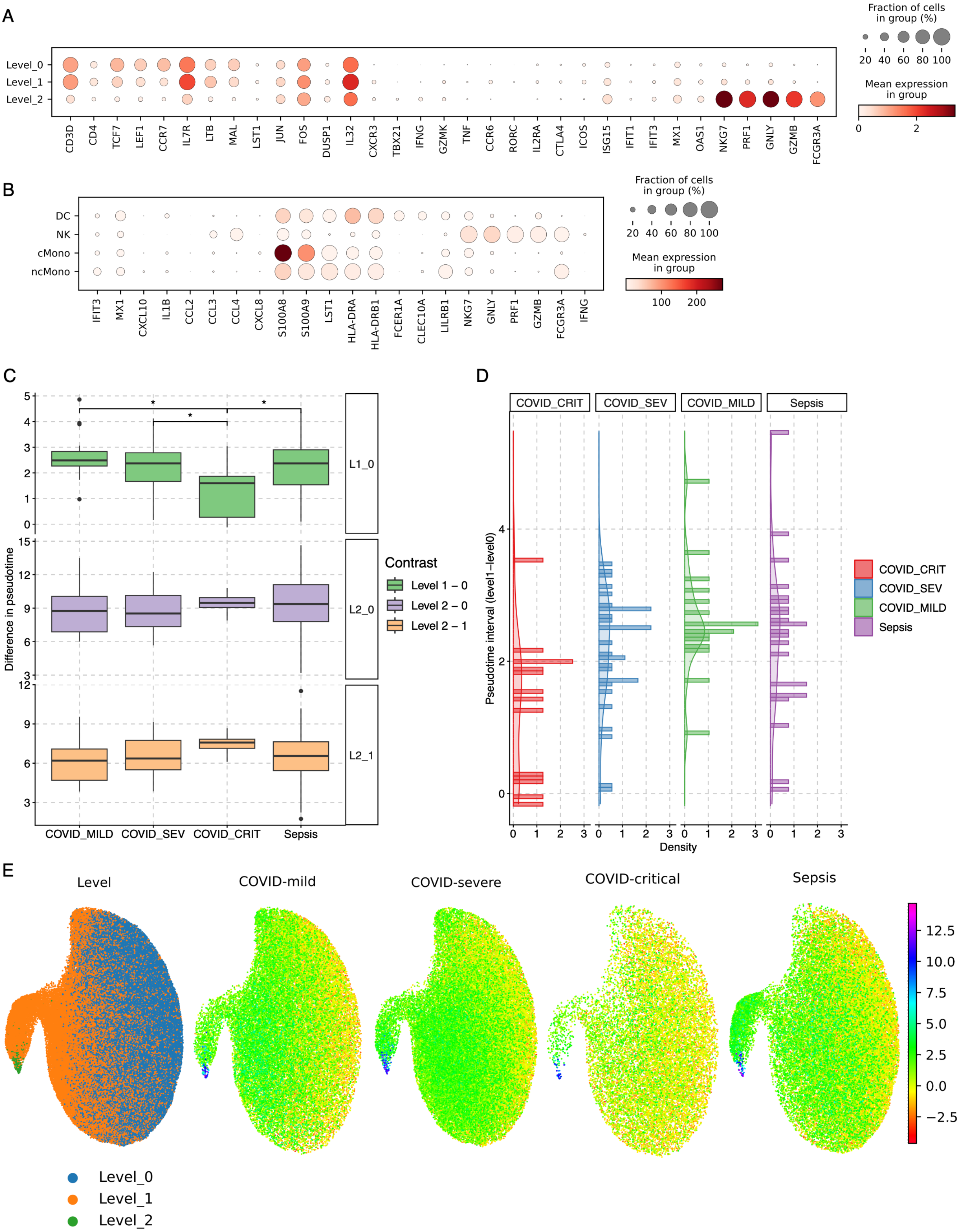
A
Fraction of cells in group (%)
20 40 60 80 100
Mean expression in group
0 2
Level_0
Level_1
Level_2
CD3D CD4 TCF7 LEF1 CCR7 IL7R LTB MAL LST1 JUN FOS DUSP1 IL32 CXCR3 TBX21 IFNG GZMK TNF CCR6 RORC IL2RA CTLA4 ICOS ISG15 IFIT1 IFIT3 MX1 OAS1 NKG7 PRF1 GNLY GZMB FCGR3A
B
Fraction of cells in group (%)
20 40 60 80 100
Mean expression in group
100 200
DC
NK
cMono
ncMono
IFIT3 MX1 CXCL10 IL1B CCL2 CCL3 CCL4 CXCL8 S100A8 S100A9 LST1 HLA-DRA HLA-DRB1 FCER1A CLEC10A LILRB1 NKG7 GNLY PRF1 GZMB FCGR3A IFNG
C
Difference in pseudotime
L1_0
L2_0
L2_1
COVID_MILD COVID_SEV COVID_CRIT Sepsis
Contrast
Level 1 – 0
Level 2 – 0
Level 2 – 1
D
COVID_CRIT COVID_SEV COVID_MILD Sepsis
Pseudotime interval (level1–level0)
Density
COVID_CRIT
COVID_SEV
COVID_MILD
Sepsis
E
Level
COVID-mild
COVID-severe
COVID-critical
Sepsis
Level_0
Level_1
Level_2

Supplementary Figure S7: Canonical markers of input data and pseudotime contrast. (A) Canonical CD4 markers across CD4 transcriptomic states, including level 1 (naive), level 2 (effector), and level 3 (terminal). (B) Canonical innate markers of microenvironmental context defined by classical monocytes, non-classical monocytes, dendritic cells, and natural killer cells. (C) Posterior median of donor-specific pseudotime coordinates across CD4 transcriptomic states across different severities of COVID-19 and sepsis. Pseudotime estimates are summarized by the median and interquartile range for each transcriptional state. * denotes statistically significant contrasts using Welch's one-way ANOV, followed by Games–Howell post hoc test for pairwise comparisons (adjusted p-value $< 0.05$). (D) Distribution of pseudotime transition intervals between level $0 \rightarrow 1$, shown as histograms with kernel density estimates. (E) Uniform Manifold Approximation and Projection (UMAP) of CD4 transcriptomic states using 500 highly variable genes, colored by estimated pseudotime coordinates.

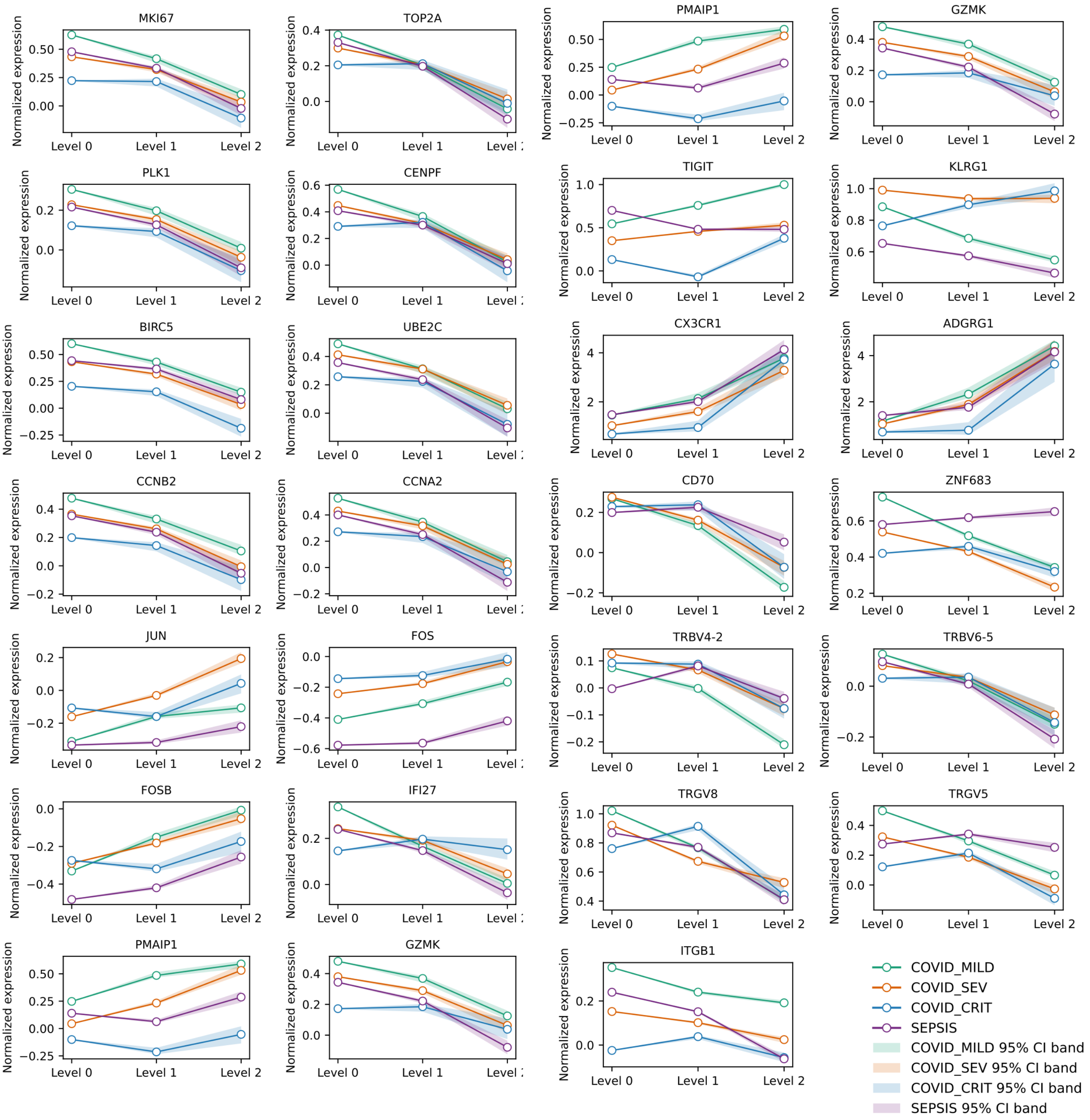


Supplementary Figure S8: Estimated denoised gene expression of pseudotime-associated genes across COVID-19 severity groups and sepsis. Shaded regions represent 95% credible intervals.

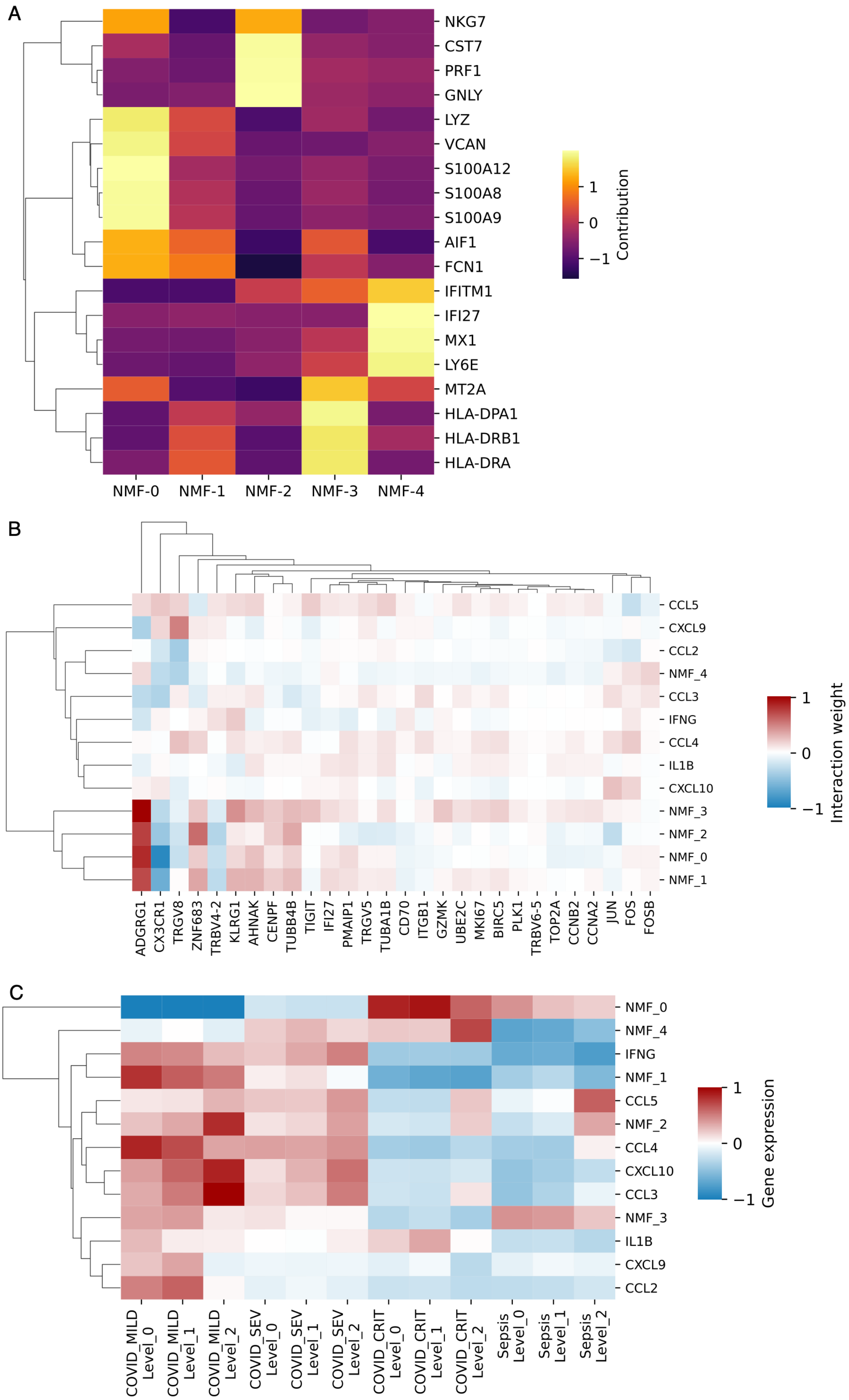
A
NKG7
CST7
PRF1
GNLY
LYZ
VCAN
S100A12
S100A8
S100A9
AIF1
FCN1
IFITM1
IFI27
MX1
LY6E
MT2A
HLA-DPA1
HLA-DRB1
HLA-DRA
NMF-0
NMF-1
NMF-2
NMF-3
NMF-4
Contribution
1
0
−1
B
CCL5
CXCL9
CCL2
NMF_4
CCL3
IFNG
CCL4
IL1B
CXCL10
NMF_3
NMF_2
NMF_0
NMF_1
ADGRG1
CX3CR1
TRGV8
ZNF683
TRBV4-2
KLRG1
AHNAK
CENPF
TUBB4B
TIGIT
IFI27
PMAIP1
TRGV5
TUBA1B
CD70
ITGB1
GZMK
UBE2C
MKI67
BIRC5
PLK1
TRBV6-5
TOP2A
CCNB2
CCNA2
JUN
FOS
FOSB
Interaction weight
1
0
−1
C
NMF_0
NMF_4
IFNG
NMF_1
CCL5
NMF_2
CCL4
CXCL10
CCL3
NMF_3
IL1B
CXCL9
CCL2
COVID_MILD Level_0
COVID_MILD Level_1
COVID_MILD Level_2
COVID_SEV Level_0
COVID_SEV Level_1
COVID_SEV Level_2
COVID_CRIT Level_0
COVID_CRIT Level_1
COVID_CRIT Level_2
Sepsis Level_0
Sepsis Level_1
Sepsis Level_2
Gene expression
1
0
−1

Supplementary Figure S9: Microenvironmental features used to model pseudotime in CD4 transcriptomic states in COVID-19 and sepsis. (A) Normalized gene weights (H matrix) for five non-negative matrix factorization (NMF) modules. Yellow indicates relatively higher gene weights within a module, and black indicates lower gene weights after normalization. (B) Estimated coefficients linking microenvironmental features to pseudotime-associated gene signatures. Color indicates the direction of the effect; white denotes non-credible interactions under the 95% credible interval. (C) NMF-derived module activity and gene expression of microenvironmental features across CD4 transcriptomic states in COVID-19 and sepsis. Red indicates a credibly positive change across lineages based on 95% credible intervals (CrIs), and blue indicates a negative change. Features with non-credible changes are shown in white.

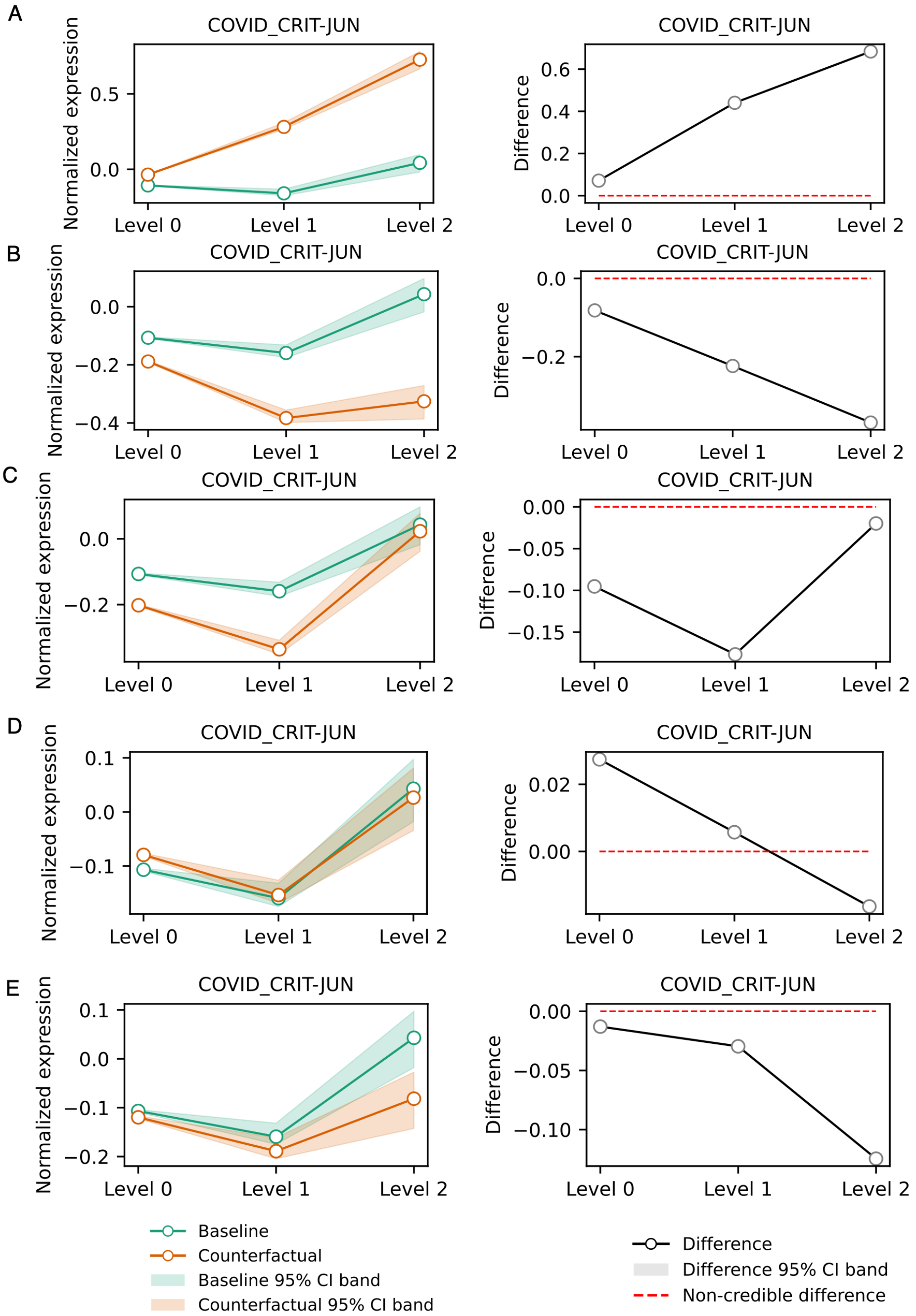


Supplementary Figure S10: Counterfactual intervention on critical COVID-19 samples by modulating microenvironmental features to the mean levels observed in mild COVID-19. Left panels show baseline and counterfactual denoised gene expression, with shaded regions indicating 95% credible intervals (CrIs). Right panels show the difference between counterfactual and baseline gene expression, with shaded regions indicating 95% CrIs. (A) Counterfactual decrease of NMF0 to the mean level observed in mild COVID-19 samples for gene JUN. (B) Counterfactual increase of NMF1 to the mean level observed in mild COVID-19 samples. (C) Counterfactual increase of NMF2 to the mean level observed in mild COVID-19 samples. (D) Counterfactual increase of NMF3 to the mean level observed in mild COVID-19 samples. (E) Counterfactual decrease of NMF4 to the mean level observed in mild COVID-19 samples.

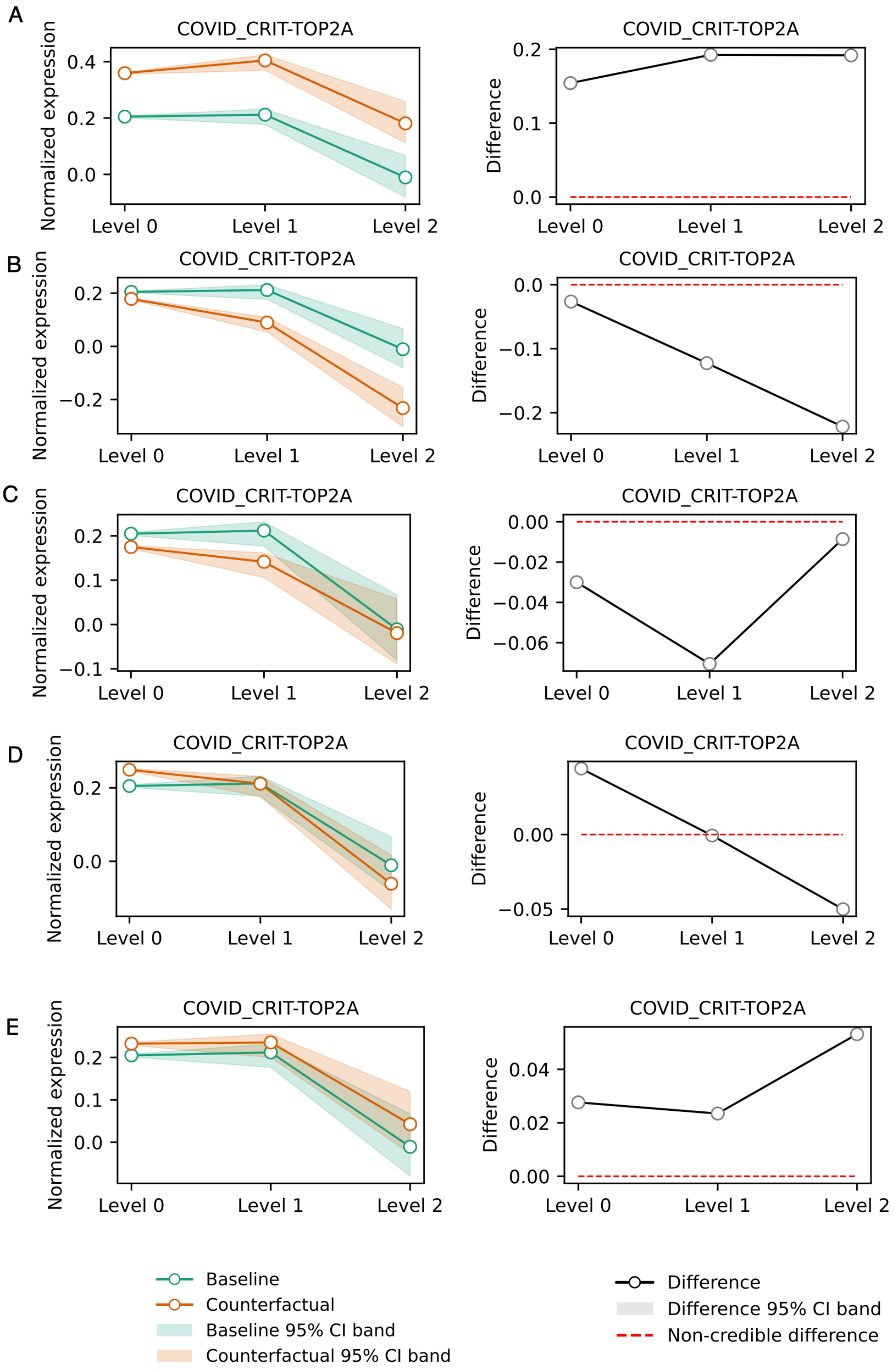


Supplementary Figure S11: Counterfactual intervention on critical COVID-19 samples by modulating microenvironmental features to the mean levels observed in mild COVID-19. Left panels show baseline and counterfactual denoised gene expression, with shaded regions indicating 95% credible intervals (CrIs). Right panels show the difference between counterfactual and baseline gene expression, with shaded regions indicating 95% CrIs. (A) Counterfactual decrease of NMF0 to the mean level observed in mild COVID-19 samples for gene TOP2A. (B) Counterfactual increase of NMF1 to the mean level observed in mild COVID-19 samples. (C) Counterfactual increase of NMF2 to the mean level observed in mild COVID-19 samples. (D) Counterfactual increase of NMF3 to the mean level observed in mild COVID-19 samples. (E) Counterfactual decrease of NMF4 to the mean level observed in mild COVID-19 samples.
